# TERRA engages MDA5 to shape the immune landscape in ALT brain tumors

**DOI:** 10.64898/2026.11.19.767868

**Authors:** Irene Pecchini, Vittorio Bontempi, Cecilia Isidori, Silvia Sorze, Alessandra Liuzzo, Inmaculada Martínez Olmo, Maria Luisa Cayuela, Toma Tebaldi, Nourhan Abdelfattah, Kyuson Yun, Eirini Trompouki, Marina Mione

## Abstract

In cancer, telomere maintenance is mediated by telomerase reactivation or by the alternative lengthening of telomere (ALT) pathway. Both mechanisms contribute to cancer cell replicative immortality, but ALT, generates telomere instability and promotes inflammation, thus contributing to cancer progression also through non-cell autonomous mechanisms. How ALT telomeres modulate inflammatory signals in cancer is not fully understood, mostly because of the lack of organismal models of ALT+ cancers. Here we used a zebrafish model of ALT+ brain tumor to demonstrate that ALT telomeres promote inflammation through increased transcription of TERRA, which engages RNA-sensing pathways through MDA5. Analysis of ALT+ brain tumors shows that MDA5-MAVS signaling results in the production of interferon responsive genes and cytokines that induce a pro-tumoral immune microenvironment, which can be prevented by abrogating TERRA signaling through *in vivo* antisense technology, or by CRISPR/Cas9 mediated knock-out of MDA5 or MAVS. Single-cell transcriptomic analysis reveals that abrogating TERRA signaling using ASOs affects both cancer cells and immune cell compartments in a long-lasting manner. These findings reveal a previously unrecognized role of TERRA in modulating immune responses in ALT+ brain tumors and highlight the potential of TERRA-targeting ASOs as a novel therapeutic strategy for ALT tumors.

## Background

Telomeres are nucleoprotein structures that protect the ends of linear chromosomes from being recognized as double-strand breaks. In somatic cells, telomeres shorten at every cell division, leading to senescence and cell-cycle arrest^1^. Telomeres protection is essential to preserve cell function and viability, however several factors can compromise telomere stability, resulting in premature aging syndromes and elevated susceptibility to cancer^2–5^. Moreover, telomere deprotection can results in chromosomal bridges, micronuclei formation^6^ and cytoplasmic chromatin fragments (CCFs) which serve as signals for pattern-recognition receptors (PRRs) and activation of innate immunity pathways^7,8^. To grow, cancer cells need to achieve replicative immortality by elongating telomeres^9^. Telomere maintenance mechanisms (TMMs) in cancer are essential to maintain genome integrity and rely either on the reactivation of the telomerase enzyme (85-90% of tumors)^10^ or on the Alternative Lengthening of Telomere mechanisms, named ALT and based on homologous recombination^11^. Due to the recombinogenic nature of this mechanism, ALT tumors are characterized by DNA damage at telomeres^12,13^, heterogeneous telomere length^14^, the formation of ALT-associated PML bodies (APBs)^15^, and elevated expression of the Telomeric Repeat-containing RNA called TERRA^16–19^. TERRA is a lncRNA conserved across different organisms, from yeast to vertebrates including higher order mammals^20–26^. TERRA in human cells is transcribed by RNA polymerase II from many, if not all, telomeres starting from the subtelomeric regions towards telomere ends^27,28^, therefore, it encompasses a 5’end containing chromosome-specific sequences and the 3’end made of a variable number of 5’-UUAGGG-3’ telomeric repetitions^24,29^. The nature of the increase in expression of TERRA in ALT cancer cells is still not completely clarified, and its role in tumors is debated. The main hypothesis is that this population of lncRNAs can favor telomere recombination by forming DNA-RNA hybrids at telomeres, known as R-loop, that induce the opening of the DNA double-stranded helix^30–32^. Apart from the context of ALT tumors, recent evidence indicates that TERRA is also involved in regulating telomerase-dependent telomere elongation by binding to telomerase RNA^33,34^. Moreover, in a non-cancerogenic condition, TERRA can trigger type-I-IFN-dependent cell death, preventing cancer initiation^35^. However, the functional relevance of these observations is still largely unexplored in a physiological context. In fact, most of the evidence linking TERRA to telomere maintenance and to the activation of innate immunity has been gathered in cell culture systems, which cannot recapitulate the complex interplay between tumor cells and the host immune microenvironment. Whether the increase in TERRA expression observed in ALT+ tumors translates into an actual pro- or anti-tumoral effect in a living organism, and whether it shapes the immune response in a way that favors or opposes tumor progression, remains unknown. The zebrafish shares with human key aspects of immune system development and functions^36–38^, as well as telomere biology and length^39^. Also, the ease of genetic manipulation allowed the generation of different transgenic tumor models^40–42^. These aspects make zebrafish a suitable model to study the role of TMM and the immune system in tumor development and progression. In particular, a previously described zebrafish glioblastoma model^43,44^, characterized by the activation of the ALT pathway, represents a suitable model to investigate the consequences of ALT on tumor progression and its immune microenvironment in a living organism. In zebrafish brain tumors, as in some human cancers, the ALT phenotype is associated with a different median survival and prognosis^12,44,45^. Given the diverse role that TERRA can play in cells, we speculated that it may be involved in determining tumor growth and aggressiveness. Glioblastoma tumor masses in humans are characterized by a strong infiltration of immune cells, in particular microglia and macrophages, usually displaying pro-tumoral functions and favoring tumor growth^46–48^. Here, we used the zebrafish ALT+ glioblastoma model to describe the role that TERRA has in shaping the immune system during tumor development through the activation of an RNA sensor pathway. This sensing results in increased infiltration by macrophages associated with increased proliferation of tumor cells. We also show that preventing the activation of RNA sensing by targeting TERRA can have a beneficial effect on tumor-bearing animals, opening the possibility to target this pathway in ALT+ tumors in order to prevent a pro-tumoral shift of the immune landscape.

## Methods

### Animal housing and line generation

Adult zebrafish (Danio rerio) were housed at the Department of Cellular, Computational and Integrative biology (CIBIO), University of Trento, and maintained in the Model Organism Facility under standard conditions (Monte Westerfield, Leonard I. Zon, H. William Detrich III. Essential Zebrafish Methods: Cell and Developmental Biology. 1st Edition). All zebrafish experimental procedures adhered to the European law on Animal Protection and were authorized under permit no. 425/2024-PR by the Italian Ministry of Health issued to M.C. Mione. Embryos were kept at 28.5°C in E3 solution. The ASOs used were Locked Nucleic Acid (LNA)-enhanced MixmeR oligonucleotides (LNA/DNA mixmers) with a fully phosphorothioate backbone (LNA™, Exiqon).

The following transgenic zebrafish lines were used in this study:

- Et(zic4:Gal4TA4, UAS:mCherry)hzm5, called Zic:Gal4^43^;
- Tg(UAS:eGFP-HRASV12G)io006, referred to as UAS:RAS^68^;
- Tg(UAS:hMDA5)^utn13^, referred to as UAS:MDA5; this work
- Tg(UAS:mmRig-I)^utn14^, referred to as UAS:Rig-I; this work.

### Transgenic line generation

To generate the tg(UAS:hMDA5) and tg(UAS:mmRig-I) lines, hMDA5 (pDONR221) or mmDdx58 (pENTR223.1, Addgene #160107) were cloned into Gateway destination vectors alongside Tol2kit plasmids (#327, #302, #395) using LR clonase, with sequences confirmed via Nanopore sequencing. One-cell stage Zic:Gal4 embryos were co-injected with 0.25 ng/μl construct plasmid and 0.25 ng/μl Tol2 transposase mRNA, then raised at 28.5 °C in E3 medium. Heart-GFP positive adults were outcrossed to Zic:Gal4 fish to establish F1 lines. Multiple independent F1 lines were screened for insertion effects; finding none, a single representative allele per transgene was selected.

### Survival curve experiment and analysis

Larvae were selected using a Leica MZ10F Stereomicroscope. The sample size per treatment group was 30 zebrafish larvae and the experiments were conducted in triplicate. The experimental groups included non-injected Zic:RAS larvae, Zic:RAS larvae injected with TERRA ASO mixmer, and Zic:RAS larvae injected with Control ASO mixmer. From 5 to 15 dpf, larvae were placed in tanks containing fish-system water and kept at 28.5°C in the Model Organism Facility. Feeding was performed daily using ZEBRAFEED <100 standardized diet for zebrafish (Sparos). The water was changed daily. Larval survival was monitored daily, and data were recorded on Microsoft Excel. GraphPad Prism (version 8.0.0) software was used to conduct statistical analyses and generate survival plots.

### Generation of crispant larvae and ASO injection

To assemble gRNA duplexes, equal volumes of 100 μM synthetic crRNA and tracrRNA (IDT) were mixed, denatured at 95 °C for 5 minutes, and cooled at room temperature for 10 minutes to yield a 50 μM (1,715 ng/μL) duplex. Ribonucleoprotein (RNP) complexes were assembled in a 5 μL reaction containing 1 μg/μL Cas9 protein, 343 ng/μL gRNA duplex (or combined multiplexed duplexes), Phenol Red tracer, and Nuclease-Free Duplex Buffer. The RNP mixture was microinjected into the yolk of one-cell stage zebrafish (Danio rerio) embryos. Two nanoliters of 2 μM TERRA ASO were injected in each 1 cell stage embryo.

### Cell line culture and transfections

Human osteosarcoma (U2OS) cells were obtained from ATCC. All cells were cultured in Dulbecco’s Modified Eagle Medium (DMEM), supplemented with 10% FBS, 1% L-Glutamine and 1% penicillin/streptomycin at 37°C in a humidified atmosphere containing 5% CO2. Cell lines were tested regularly for mycoplasma contamination by the CellTech CIBIO facility. ASO transfections were performed in U2OS cells using Oligofectamine Transfection Reagent (Invitrogen) according to the manufacturer’s protocol. ASOs were transfected at the final concentration of 1 μM.

### RNA extraction and Northern Blot

Total RNA was extracted using TripleXtractor Reagent (Grisp) from homogenized adult zebrafish brains (N=1/sample), pooled larvae (15-20/sample), or direct cell lysates, and quantified via NanoDrop Lite (Thermo Fisher Scientific). For northern blotting, 1 μg total RNA was glyoxal-denatured (50 °C, 30 min), resolved on a 0.9-1.0% agarose/MOPS gel (80-85 V, 2-2.5 h), and transferred overnight to a membrane by downward capillary blotting in 10X SSC followed by UV crosslinking. Membranes were pre-incubated in 50 mM Tris-HCl (pH 8.0, 45 °C, 20 min), hybridized in ULTRAhyb-Oligo Buffer (42 °C, 1 h), and incubated overnight with a biotinylated probe. After sequential SSC/SDS washes at 42 °C, signals were developed using Streptavidin-HRP with chemiluminescent substrate and imaged on a ChemiDoc system.

### RNA immunoprecipitation (RIP)

Adapted from Keene et al. (2006)^69^, RNA immunoprecipitation (RIP) was performed using confluent U2OS cells from two to five 10 cm plates per condition. Cells were washed with 1X PBS and scraped on ice in Polysome lysis buffer containing protease and RNase inhibitors. Lysates were cleared by centrifugation (12,000xg, 20 min, 4 °C) and pre-cleared with uncoated Dynabeads Protein G (2.5h, 4 °C). In parallel, 25 μL Dynabeads Protein G was coated with 3 μg anti-MDA5 (Proteintech, 21775-1-AP) or control mouse IgG (Millipore) in NT2 buffer with inhibitors for 2.5h at 4 °C. Pre-cleared lysates (2-5 mg protein; quantified via BCA) were incubated overnight at 4 °C with antibody-bead complexes; 200-600 μg protein was reserved as INPUT. Beads were washed four times with NT2 buffer containing inhibitors and 0.01% urea at 4 °C, then resuspended in inhibitor-free NT2. Bound complexes and INPUT samples were split for Western blot analysis and TripleXtractor-based RNA isolation for RT-qPCR.

### Retro transcription and qPCR analysis

For each RIP experiment, a total of 1-2 μg of extracted RNA was subjected to genomic DNA (gDNA) removal and reverse transcription using the ReverTra AceTM qPCR RT Master Mix with gDNA remover (Toyobo) according to the manufacturer’s protocol. Briefly, RNA samples were denatured by incubation for 5 min at 65°C and immediately placed on ice. The 4X DN Master Mix (containing gDNA remover) was added, and samples were incubated at 37°C for 5 min to digest residual gDNA. For no-reverse transcriptase (-RT) controls, 3 μl was taken from each sample and combined with 5X RT Master Mix II no RT-Control. The remaining volume was reverse-transcribed by adding 5X RT Master Mix II, supplemented with 0.2 μL of COMMON TERRA antisense RT primer (10 μM). Both reverse transcription (RT) and -RT reactions were incubated under the following conditions: 37°C for 15 min, 50°C for 5 min, and 98°C for 5 min. Finally, the resulting cDNA samples were diluted 1:3 in nuclease-free H2O for downstream qPCR analysis. Quantitative real-time polymerase chain reaction (qPCR) was performed on cDNA using a CFX96 Real-Time Systems (Bio-Rad) machine and 2X qPCRBIO SyGreen® Blue Mix No-ROX (PCR Biosystems). Primers were added to the qPCR reaction mix at a final concentration of 0.4 μM. cDNA was amplified in triplicate in a total reaction volume of 10 μl (8 μl of reaction mix and 2 μl of cDNA). Primers used in this study are listed at the end of this chapter (paragraph 5.12). A β-Actin and 18S rRNA were used as housekeeping genes for human cells, RPS11 was used for zebrafish. Gene expression levels were normalized to the respective housekeeping genes and analyzed using Bio-Rad CFX Manager software (version 3.1).

### Western blot

Extracted proteins were added with 4X Loading buffer and β-Mercaptoethanol as reducing agent, and then denatured by boiling at 95°C for 5 min. For RIP experiments, a magnetic rack was used to prevent magnetic beads loading into the gel. Protein were resolved by sodium dodecyl sulfate-polyacrylamide gel electrophoresis (SDS-PAGE) and blotted on polyvinylidene difluoride (PVDF) membranes (Bio-Rad). The PM2610 ExcelBand™ (SMOBIO) was used as a protein size marker. SDS-PAGE was initially run at 60 V for 30 min, and subsequently at 120 V for protein separation. Transfer on the PVDF membrane was performed at 300 mA for 2 hours. Membranes were blocked with 5% nonfat dried milk powder (PanReac AppliChem) in TBS-T for 1 hour at room temperature. The membranes were then incubated overnight at 4°C with primary antibodies diluted in 1% milk in TBS-T according to the data sheets. Membranes were washed four times in PBS-T and incubated with an HRP-conjugated secondary antibody diluted in 1% milk in TBS-T for 1 hour at room temperature. After washing, the membranes were incubated with Trident Plus Western HRP Substrate (GeneTex). Chemiluminescent bands were detected using a ChemiDoc™ Imaging System (Bio-Rad). Western blotting was performed using the primary antibodies and HRP-conjugated secondary antibodies listed in Table 5.3.

### Histology and RNA-FISH

Fresh tissue sections were fixed in 4% PFA for 15 min a RT, and washed twice in PBST (0.05% Tween-20). All tissue sections were permeabilized in 70% ethanol at 4°C O/N. For RNase control samples, slides were washed twice with 2X SSC and incubated 4 mg/mL RNaseA in 2X SSC for 3 hours at 37°C in a humidified chamber prior to hybridization. All samples were washed twice with 2X SSC and equilibrated in wash buffer (Table 5.7) for 3-5 min at RT. Tissues were then covered with 150 μL of hybridization buffer (Table 5.8) and incubated overnight at 37°C using Parafilm coverslips in a humidified chamber. Post-hybridization, slides were rinsed in wash buffer for 5 min at RT, incubated in wash buffer at 37°C for min, and rinsed again for 5 min at RT. Nuclei were counterstained with 2g/mL DAPI in 2X SSC for 5 min, followed by a single wash in 2X SSC-T (0.1% Tween-20). Sections were mounted in antifade mounting medium and sealed with nail polish before imaging.

### Wholemount immunofluorescence on brains

Whole-mount was performed on brains extracted from fish larvae at 5 dpf. Briefly, 5 dpf-zebrafish larvae were fixed in 4% PFA in PBS O/N at 4°C or 3 hours at RT. The samples were then subjected to multiple washes with PBS 1X, with each wash lasting a minimum of 1 hour. Brains were extracted from the larvae using a Sylgard plate and forceps. Extracted brains were permeabilized and blocked in whole-mount blocking buffer (Table 5.4) for 1 hour at room temperature under agitation. Brains were then incubated O/N at 4°C under agitation with primary antibodies (Table 5.5) in blocking buffer. Multiple washes of at least 1 hour each were performed in PBS 0.05% Triton X-100. The brains were then incubated O/N at 4°C or 2 hours at room temperature with fluorescent secondary antibody in blocking buffer and then subjected to multiple washes in PBS 0.05% Triton X-100. Entire brains were mounted on antifade solution (1% n-propylgelat, 90% glycerol, and 10% PBS 10X) and acquired with a Nikon Eclipse Ti2 inverted confocal microscope: rabbit anti-L-plastin (1:1000) or rabbit anti-pHH3 (Upstate 06-570, 1:1000).

### PLA

Proximity Ligation Assay (PLA) was performed following the Duolink® PLA Fluorescence Protocol provided by Merck (Sigma Aldrich). Cellular samples on coverslips were pre-treated with respect to fixation and permeabilization. Briefly, HeLa and U2OS cells, transfected or not with the TERRA-targeting ASO TeloG, were fixed in cold 4% PFA in PBS for 5-10 min and then permeabilized in 1% Saponin in PBS for 1 hour at room temperature on a shaking platform. Three washes of 5 min each were performed in 1X PBS at room temperature. After the last wash, cells were blocked in Blocking Buffer solution (Table 5.6) for 3 hours at 4°C under agitation. Meanwhile, 100 nM TERRA antisense PLUS probe (Metabion) and Scrambled (Scr) PLUS probe (Eurofins Genomics) in blocking buffer solutions were denatured at 70°C for 3 min. After 3 hours of incubation, the blocking buffer was removed and replaced with 100 μL of PLUS probes (TERRA or Scr). Cells were incubated O/N at 4°C on a shaking platform. The following day, cells were washed in 1X PBS and blocked in 0.1% PBS-Tween® 20 with 1% BSA and 20 μg/ml sssDNA for 1 hours 30 min at room temperature on a shaking platform. Washes in 1X PBS and 2X SSC with 0,1% Tween® 20 were performed. Each coverslip was then incubated with rabbit polyclonal anti-MDA5 primary Ab (1:100, Proteintech, 21775-1 AP) in PBS-T 0,1% for 1 hour at room temperature in a humidified chamber. Meanwhile, the PLA probe solution was prepared by diluting the Duolink® In Situ PLA® Probe Anti-Rabbit MINUS at a 1:5 ratio into PBS-T 0,1% containing 20 μg/ml sssDNA and allowing the mixture to sit for 20 min at room temperature. All the following incubations were performed in a humidity chamber. After 1X PBS wash, coverslips were incubated with the probe solution for 1 hour at 37°C. Meanwhile, the 5X Duolink® Ligation buffer was diluted at a 1:5 ratio in Ultra-pure H2O and, after three washes in 1X Wash Buffer A, cells were incubated in 1X Ligation buffer added with Ligase enzyme (1 U/μl) at a 1:40 dilution for 30 min at 37°C. The 5X Duolink® Amplification buffer was then diluted at a 1:5 ratio in Ultra-pure H2O and added with Polymerase enzyme (10 U/μL) at a 1:80 dilution. Following the other three washes in Wash Buffer A, cells were incubated in the amplification solution for 100 minutes at 37°C. Since the Amplification buffer is light sensitive, all solutions containing the buffer were protected from light. Finally, cells were washed twice in 1X Wash Buffer B for 10 minutes at room temperature and then once in 0.01X Wash Buffer B with Hoechst 33342 (1:10000, Invitrogen). Coverslips were mounted on frosted microscope slides (Trajan) using a minimal volume of anti-fade solution and sealed with clear nail polish. Microscope slides were then acquired with a Nikon Eclipse Ti2 confocal microscope or stored in the dark at 4°C.

### PLA on zebrafish tissue, sample preparation

Brain biopsies were enzymatically dissociated incubating each sample in a solution of DMEM/F12 (Thermo Fisher Scientific) added with Liberase (1:100, Roche) and DNase I (1:500, Sigma-Aldrich) at 36°C for 15 min. Additional mechanical disintegration was performed by gentle pipetting up-and-down every 3 min during incubation. After dissociation, to neutralize the enzymatic reaction and facilitate pellet individuation DMEM/F12 supplemented with 10% FBS and 1 μL/mL methylene blue (Thermo Fisher Scientific) was added. Cell suspensions were centrifuged at 1.000 x g for 4 min at 4°C, ad supernatant was completely removed. The resulting pellets were snap-frozen in liquid nitrogen and subsequently transferred in an Optimal Cutting Temperature (OCT) compound embedding mold. The OCT block was stored at -80°C until cryosectioning. Sections were cut at a thickness of 14 μm.

### Imaging and image processing

Images where processed with Fiji and represented as maximum projection. Images of adult brains (Supplementary Fig. 1J-L) where processed and analyzed with Imaris.

### Single cell RNA seq

Biopsies from adult zebrafish bearing the tumors were collected using a scalpel and immediately placed into cold PBS droplets. Samples were kept on ice for 5 min to remove red blood cells. After centrifugation the pellet were resuspended in fresh medium and filtered through a 40 μm filter to remove aggregates. After filtration, it was centrifuged at 200 x g for 4 min, supernatant was discarded and pellet resuspended in 1000 μl 1X PBS with 10% FBS. Cells were subjected to a debris removal step using the Debris Removal Solution (Milteny 130-109-398), following the manufacturer’s instructions. Cell viability was assessed with a hemocytometer and trypan blue, together with FACS (BD FACS Symphony A1). Cells were loaded into the 10x Chromium Controller. GEM generation and barcoding, GEM-RT, amplification, and library construction were performed following the manufacturer’s instructions (Chromium Next GEM Single cell 3 or 5’ Reagent kits v3.1-Dual Index, 10x Genomics). One biological replicate of Zic:RAS was instead processed using the Illumina Single cell 3’ RNA prep following the manufacturer’s instructions. The libraries were then sequenced on a NOVASEQ6000 with a 28-base read 1 and a 90-base read 2, 812 aiming for 25.000 reads per cell (45-base read 1 and 72-base read 2 for the illumina libraries). Raw sequencing reads were aligned to a custom zebrafish reference using Cell Ranger v7.1.0, with default parameters. The custom zebrafish reference was created using Cell Ranger v.7.1.0 by adding the sequence of the transgene eGFP-HRASV12G to the GRCz11 reference genome and Lawson Lab Transcriptome Annotation188, V4.3.2. Ambient RNA was removed using Soupx 1.6.2189, and high-quality cells containing <10% mitochondrial transcripts and >200 total transcripts were retained for further analysis. Genes expressed in fewer than three cells were discarded. Doublets were removed using db_scDblFinder from the SCP R package (version 0.5.6)^67^, with an expected doublet rate of 1% per 1.000 cells. Counts were normalised to the library size, scaled across all genes, and principal components analysis (PCA) was performed on the 2,000 most variable genes. Samples were integrated using Harmony v1.2.3 with the RunHarmony R function191. Clusters were defined using Seurat 5.1192. Annotation was performed manually based on literature mining. The cell types identified in our analysis expressed the following marker genes: cancer cells (expressing *eGFP-HRASv12g, fabp7a, slc1a2b, mdkb, prss35* and *gpr37l1a*, corresponding to clusters C03, C24); NK/T cells (expressing the T cell receptor beta orthologs *FP236356.1, cd4-1, cd8a, cd8b, nkl.3, nkl.4, ccl38.6, il2rb* and *gzm3.3*, corresponding to clusters C08); endothelial cells (expressing *pecam1, tek, krt8 and epas1b*, corresponding to cluster C05, C17); pericytes (expressing *acta2* and *pdgfrb*, corresponding cluster C12); macrophages/microglial cells (expressing *csf1rb, csf1ra, spi1a, lgals3bpb, mpeg1.1, marco* and *c1qb*, corresponding to clusters C04, C18, C31); astrocytes (expressing *gfap* and *ptprz1a*, corresponding to cluster C02, C13); oligodendrocyte (expressing *mbpa, mbpb* and *ppp1r1c*, corresponding to C07); oligodendrocyte progenitor cell (expressing *olig2, mpz* and *plp1b*, corresponding to C25); radial glia cell (expressing *fabp7a, slc1a2b* and *mdkb*, corresponding to C01, C11, C15, C22, C26); quiescent radial glia cells (expressing *slc1a3b, s100b, glula, fabp7a, slc1a2b* and *mdkb*, corresponding to C14); fibroblasts (expressing *col1a1a, fn1a* and *lamc3*, corresponding to clusters C16); lymphatic endothelial cell (expressing *stab1, mrc1a* and *lyve1b*, corresponding to C21); B cells (expressing *blnk* and *dapp1*, corresponding to clusters C29); T-helper 2 cells (expressing *il4, il13, cd247* and *FP236356.1*, corresponding to cluster C27); neurons (expressing *snap25a, egr4, kiss1* and *neurod1*, corresponding to C06, C10, C19, C23, C28, C30); dendritic cells (expressing *xcr1a.1, siglec15l* and *flt3*, corresponding to C20); neural progenitor proliferative cell (expressing *neurod1, mki67* and *pcna*, corresponding to to C09). Cells annotated as cancer cells, or immune cells, were subset from the original dataset and analyzed as described above (normalization, scaling, dimensionality reduction and integration). De novo clustering was done to identify subpopulations of the cells of interest. Differential expression analysis to find significant genes among two given groups was carried out using the pseudobulk approach. Differential expression was conducted using Deseq2 with Wald test for statistics. Gene set enrichment analysis (GSEA) was carried out on differentially expressed genes between groups of interest using the wilcoxauc function (presto R package v1.0.0). Differentially expressed genes were filtered for adjusted p-value < 0.05 and logFC > 0.1. GSEA was conducted with the fgsea function (fgsea 1.32.4 R package) with the Hallmark AND Reactome pathways gene sets obtained from MsigDB. Normalized enriched scores (NES) were calculated, and results were visualized using heatmaps (pheatmap v1.0.12). For each group, the top upregulated and downregulated pathways were chosen for visualization.

### Statistical analysis

Details about the statistical analysis for each figure are provided in the figure legends. Significance is indicated as follows: ∗: p < 0.05, ∗∗: p < 0.01, ∗∗∗: p < 0.001, ∗∗∗∗: p < 0.0001, ns: not significant. GraphPad Prism 8 was used for statistical analyses.

### List and description of oligonucleotides used in the study

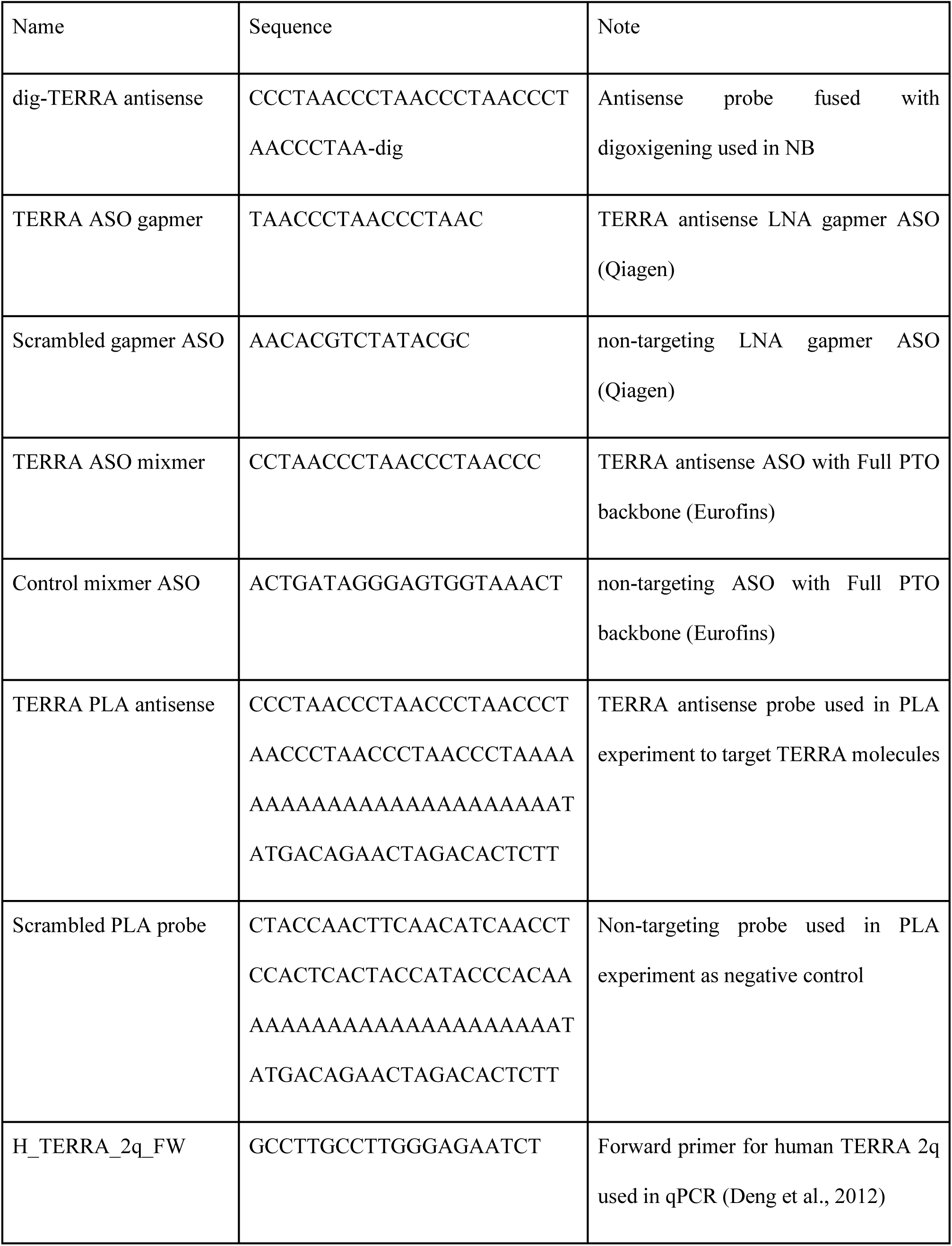
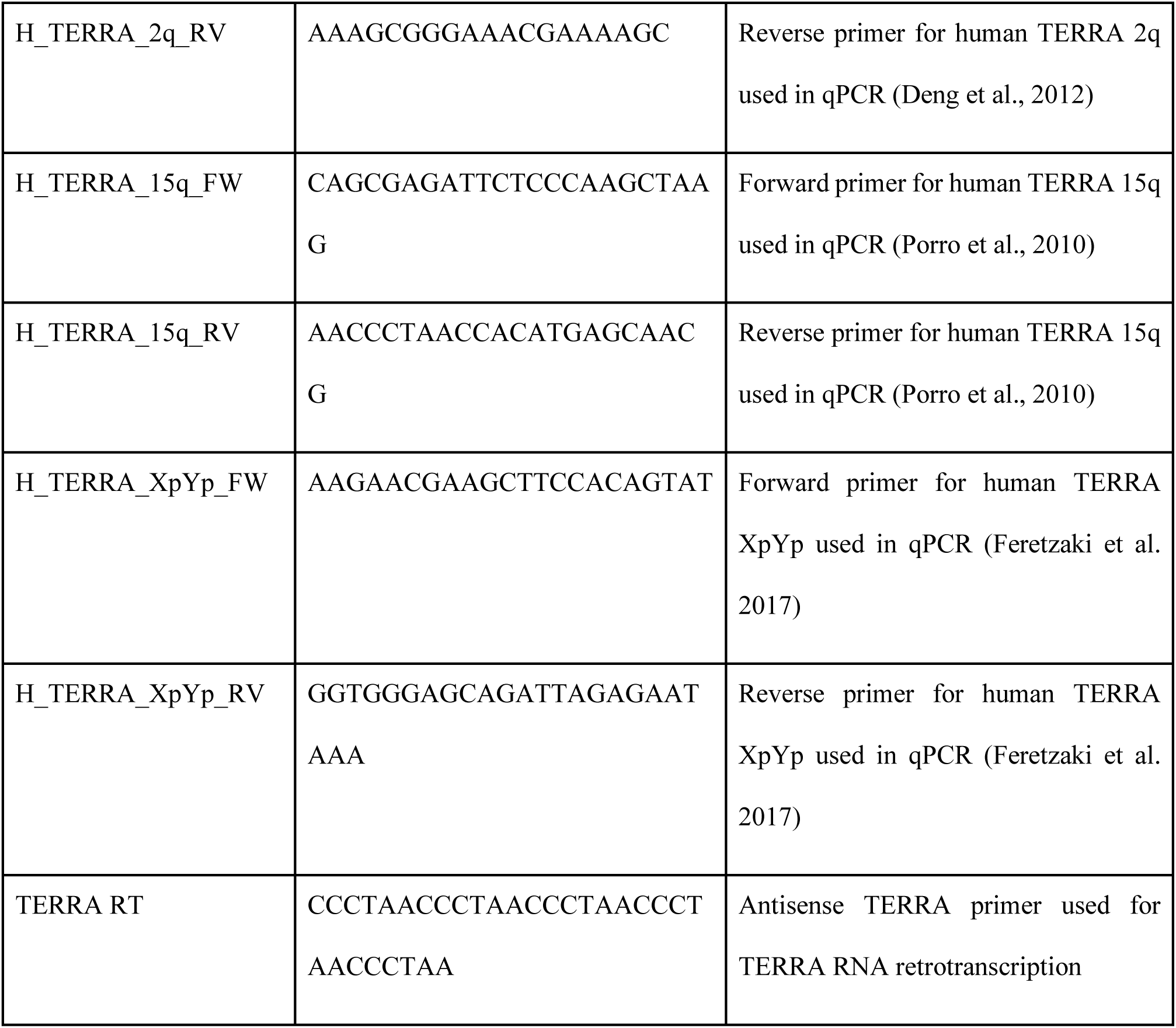

## Results

### 1. Increase of TERRA expression in ALT+ brain tumors correlates with increased inflammatory responses

Our previous study^44^ revealed that brain tumors developing in the zebrafish Zic:RAS model were not only ALT+, but also exhibited a striking upregulation of TERRA, detected already at 9 days post fertilization (dpf). This early overexpression suggests that, in this model, increased TERRA expression was the earliest molecular indicator that Telomere Maintenance Mechanisms (TMM) were leaning towards ALT. To test this hypothesis, we studied TERRA expression profile in Zic:RAS larvae (5 dpf) and adult brains, using Northern blot, RNA-FISH and qPCR. Northern blot analysis demonstrated that TERRA expression was higher in Zic:RAS larvae compared to Zic:Gal4 controls (Figure 1A-B), and the increase of TERRA expression was present, although less pronounced, also in adult tumor brain tissue (Supplementary 1A-B). Qualitative assessment of TERRA expression through RNA-FISH revealed a marked accumulation of TERRA foci in both larval (Figure 1D-F) and adult brains (Supplementary 1D-F) from Zic:RAS fish. Notably, the tumors exhibited significant cell-to-cell heterogeneity in foci distribution (Figure 1F, Supplementary 1F). Given a recent report that elevated TERRA can trigger innate immune pathways via MAVS^35^, we reasoned that these tumors might harbor a distinct inflammatory microenvironment. To test this, we compared Zic:RAS transcriptomic profiles against wild-type fish at both adult and larval stages. Our analysis showed increased expression of inflammatory genes, including cytokines (Figure 1G, Supplementary 1G), NF-κB targets genes and type I interferon-responsive genes (Figure 1H, Supplementary 1H) along with nucleic acid sensing components (Figure 1I, Supplementary 1I) in Zic:RAS tumors compared to control at both stages analyzed (Supplementary table 1). To assess whether an activation of nucleic acid sensing also occurs in ALT+ human gliomas, we interrogated a pediatric glioma dataset^49^ (see M.M.) and found that most of the genes found increased in Zic:RAS ALT+ tumors were overexpressed in pediatric diffuse gliomas (DG) carrying histone3 G34 mutations, a well-established ALT+ pediatric HGG compared to H3 WT gliomas and to HGG carrying H3 K27M mutations, a Telomerase-dependent pediatric GBM^50^ (Figure 1J, Supplementary table 2). This confirmed a trend for the increase of nucleic acid sensing in ALT+ tumors both in developing zebrafish and in pediatric HGG. GBM in human are known to be strongly infiltrated by immune cells, among which macrophages and microglia^51^. Interestingly, in zebrafish the pro-inflammatory transcriptome correlated with changes in the immune tumor microenvironment (TME). Whole-mount immunofluorescence confirmed a robust infiltration of L-plastin+ immune cells in Zic:RAS brains, clearly visible already at 5 dpf (Figure 1K-M) and maintained in adult tumors (Supplementary Fig. 1J-L). Taken together, these data indicate that genes of the RNA-sensing cascade are overexpressed in zebrafish brain tumors, and this may be correlated with the increased expression of TERRA in these tumors.

**Figure 1:**
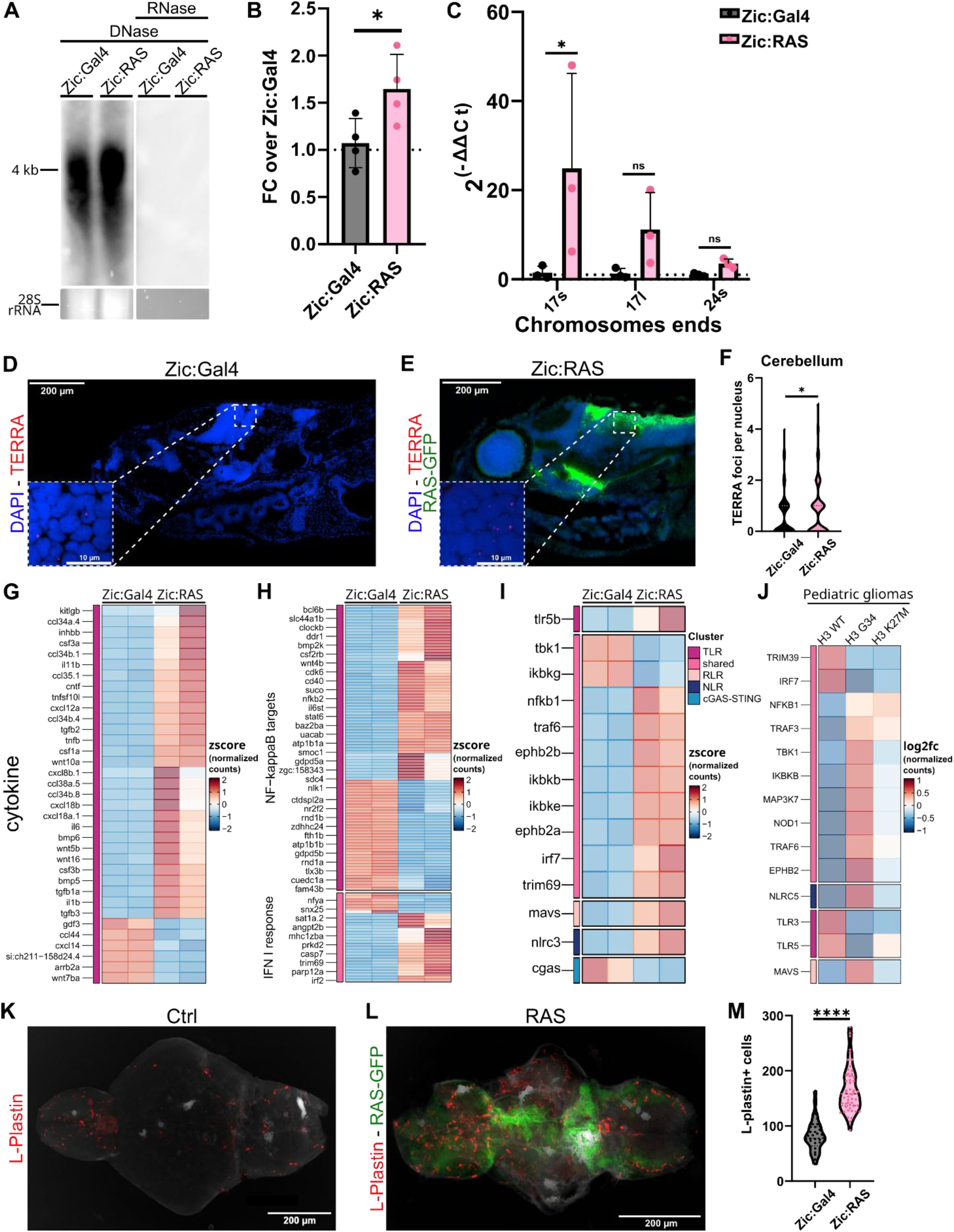
Increase of TERRA expression in ALT+ brain tumors correlates with increased inflammatory responses. (A) Representative Northern blot showing TERRA levels in Zic:Gal4 (control) versus Zic:RAS 5 dpf larvae (28s rRNA signal on gel before transfer used as loading control, shown below). (B) Quantification of relative signal intensity normalized over control. (Statistics: unpaired T-test, n=4, *p<=0.05). (C) qPCR analysis of TERRA expressed from telomeres 17s, 17l and 24s in 5 dpf larvae. Quantity is expressed as fold change of Zic:RAS over Zic:Gal4, normalized over HK gene. (Statistics: 2way Anova with Sidak’s multiple comparisons test, n=3, *p<=0.05, ^ns^p>0.05). (D, E) RNA fish performed on Zic:Gal4 (D) or Zic:RAS (E) 5dpf larvae highlighting cell nuclei (blue), RAS-GFP (green) expression and TERRA foci (red). (F) Quantification of TERRA foci per nucleus in the cerebellum of Zic:Gal4 and Zic:RAS 5dpf larvae (Statistics: Mann-Whitney test, Zic:Gal4 n=136, Zic:RAS n=131, *p<=0.05). (G, H, I) Heatmap showing the z-scores of the expression of cytokines (G), NF-kB targets, INF-I response (H) and RNA-sensing genes (I) in two pool of Zic:Gal4 or Zic:RAS 5dpf larvae. (J) Heatmap showing log2fc of the expression of RNA-sensing genes in pediatric HGG characterized by the mutational state of H3 gene: H3 WT, H3 G34 and H3 K27M). (K, L) Representative Fiji-processed max-projection WM-IF images of 5 dpf larval brains of Zic:Gal4 (J) and Zic:RAS (K), showing L-plastin staining (red) and RAS-GFP expression (green). Scale bar: 200 μm. (M) Quantification of the number of L-plastin+ cells per brain (Statistics: Mann-Whitney test, Zic:Gal4 n=36, Zic:RAS n=53, ****p<=0.0001).

### 2. Genetic manipulation of the RIG-I-like-Receptor (RLR) pathway alters immune infiltration in Zic:RAS ALT+ tumors

The primary receptor driving TERRA detection in humans is ZBP1^35^. However, the gene named *zbp1* (ZDB-GENE-060929-1258^52^) in zebrafish codes for insulin-like growth factor 2 mRNA binding protein, a protein previously known as ZBP1 in human, that has no ability to recognize left-handed Z-DNA or RNA structures^53,54^. In addition we performed, ortholog search using DIOPT (DRSC integrative ortholog prediction tool, version 10)^55^, but failed to identify a zebrafish ortholog of ZBP1. Therefore, we hypothesized that other RNA sensors were potentially interacting with TERRA in the zebrafish tumor context. Given the sequence, structure and cellular localization of TERRA^56–59^, and the receptors found overexpressed in the transcriptomic analysis (figure 1I, supplementary Fig. 1I), we hypothesized that the RLR pathway and its receptors RIG-I and MDA5 may be involved in TERRA sensing. To establish a functional link between RLR pathway activation and elevated TERRA expression, we investigated whether manipulating specific RIG-I-like receptors would affect tumor cells and immune cells infiltrating the tumor. We generated transgenic zebrafish lines overexpressing either human MDA5 (*hsaMDA5*, designated Zic-MDA5)^utn13^ or mouse RIG-I (*mmRig-I*, designated Zic-RigI)^utn14^, the two primary sensors within the RLR pathway, under the control of Zic:Gal4 through the UAS promoter (Figure 2A, Supplementary Fig. 2A-C). Our rationale was that, given the overexpression of TERRA in our Zic:RAS tumors, a low expression of RNA sensors could represent a limiting factor to understand the effects of this interactions, and so expanding the pool of available receptors would amplify downstream signals and immune cell recruitment when the appropriate sensor driving the response was overexpressed. Intriguingly, overexpression of *hsaMDA5*, but not *mmRig-I*, in tumor cells boosted immune cell infiltration within our tumor model (Figure 2C-F). On the contrary, the expression of *hsaMDA5*, as well as *mmRIG-I*, in a wild-type background (lacking tumors) resulted in a reduction in L-plastin+ cells infiltrating the brain (Supplementary Fig. 2D-G). This indicates that receptor abundance alone is not triggering the cascade involved in immune cells recruitment, but that specific pathogenic RNA ligands, such as TERRA, are present in tumor cells to license MDA5 activation. Similar results emerged when evaluating tumor cell proliferation via pHH3 immunostaining: RAS-Rig-I larvae did not show a difference in the number of pHH3+ cells in the brain, while RAS-MDA5 larvae exhibited a significant increase in pHH3+ mitotic cells compared to the baseline RAS tumor model (Figure 2G-J). Consequently, MDA5 emerged from this transgenic screen as the prime candidate for sensing tumor-derived immunogenic RNAs, like TERRA. To validate the requirement for MDA5 and its downstream signaling during brain tumorigenesis, we employed CRISPR/Cas9 technology to generate crispant larvae deficient in either MDA5 or the downstream effector MAVS (Figure 2B, Supplementary Fig. 2H). Knocking down one or the other component in RAS larvae led to a striking depletion of infiltrating L-plastin+ immune cells compared to unmodified RAS counterparts (Figure 2K-N). Furthermore, in line with the data obtained with the overexpression, this blunted inflammatory response correlated with a reduction in tumor cell proliferation, as assessed by pHH3 immunostaining (Figure 2O-R). Collectively, these genetic loss-of-function and gain-of-function experiments demonstrate that the RLR pathway, primarily through the MDA5/MAVS axis, is a key regulator of both immune cell recruitment and cell proliferation activity during early tumor development.

**Figure 2:**
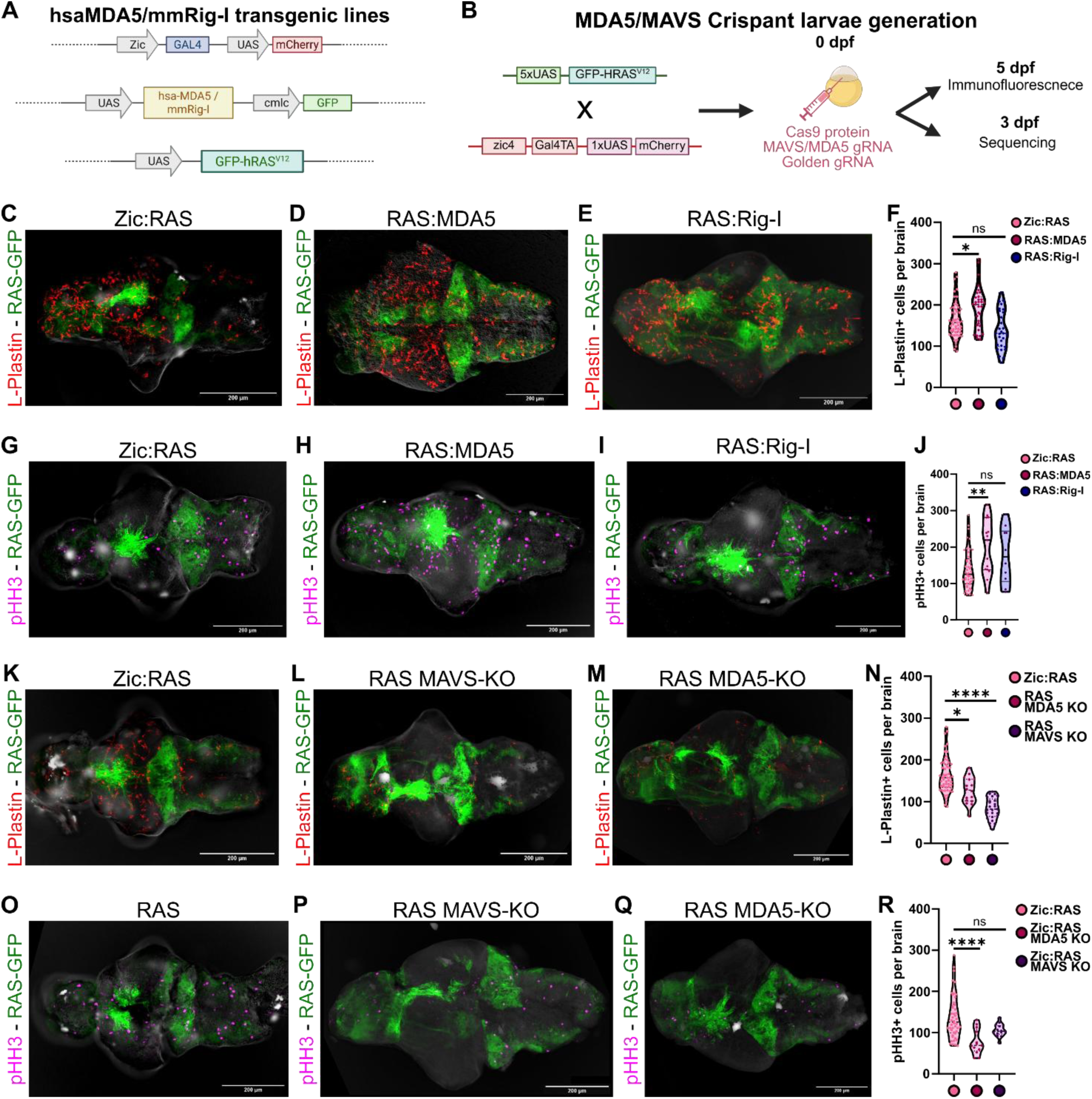
Genetic manipulation of the RIG-I-like-Receptor (RLR) pathway alters immune infiltration in Zic:RAS ALT+ tumors. (A) Schematic representation of the Gal4/UAS construct designs used to establish transgenic lines expressing hsa-MDA5 or mmRig-I along with oncogenic driver GFP-RASV12, and fluorophore marker (cmlc-GFP, UAS-mCherry). (B) Schematic representation of the protocol used to obtain and analyze crispant larvae (created with Biorender.com). (C-E) Representative WM-IF images of 5 dpf larval brains of Zic:RAS (C), RAS:MDA5 (D) and RAS:Rig-I (E), showing L-plastin staining (red) and RAS-GFP expression (green). Scale bar: 200 μm. (F) Quantification of the number of L-plastin+ cells per brain (Statistics: Kruskal-Wallis test with Dunn’s multiple comparisons test, Zic:RAS n=55; RAS:MDA5 n=33; RAS:RIG-I n=29; *p<=0.05). (G-I) Representative WM-IF images of 5 dpf larval brains of Zic:RAS (G), RAS:MDA5 (H) and RAS:Rig-I (I), showing pHH3 staining (magenta) and RAS-GFP expression (green). Scale bar: 200 μm. (J) Quantification of the number of pHH3 cells per brain (Statistics: Kruskal-Wallis test with Dunn’s multiple comparisons test, Zic:RAS n=46; RAS:MDA5 n=15; RAS:RIG-I n=10; **p<=0.01). (K-M) Representative WM-IF images of 5 dpf larval brains of Zic:RAS (K), RAS MAVS-KO (L) and RAS MDA5-KO (M), showing L-plastin staining (red) and RAS-GFP expression (green). Scale bar: 200 μm. (N) Quantification of the number of L-plastin cells per brain (Statistics: Kruskal-Wallis test with Dunn’s multiple comparisons test, Zic:RAS n=55; RAS MAVS-KO n=25; RAS MDA5-KO n=16; ****p<=0.0001; *p<=0.05). (O-Q) Representative WM-IF images of 5 dpf larval brains of Zic:RAS (O), RAS MAVS-KO (P) and RAS MDA5-KO (Q), showing pHH3 staining (magenta) and RAS-GFP expression (green). Scale bar: 200 μm. (R) Quantification of the number of pHH3 cells per brain (Statistics: Kruskal-Wallis test with Dunn’s multiple comparisons test, Zic:RAS n=46; RAS MAVS-KO n=14; RAS MDA5-KO n=13; ****p<=0.0001; ^ns^p>0.05).

### 3. Increased TERRA engages the RLR pathway to induce inflammation

Given that the RNA sensor MDA5 seems to modulate both the inflammatory landscape and cell proliferation in TERRA-overexpressing zebrafish brain tumors, we reasoned that TERRA itself might serve as a primary ligand triggering the MDA5 signaling cascade. To validate this hypothesis, we used U2-OS cells, a well-established ALT+ osteosarcoma cells model characterized by robust endogenous TERRA expression^14^. We investigated the physical interaction between TERRA and MDA5 using both RNA immunoprecipitation (RIP) and RNA-protein proximity ligation assays (PLA). Immunoprecipitation of endogenous MDA5 using specific antibodies (Figure 3A) successfully retrieved a significantly higher fraction of TERRA transcripts compared to the IgG negative control, as quantified by qPCR (Figure 3B). To validate these biochemical data and resolve the subcellular spatial dynamics of this interaction, we employed an RNA-protein PLA imaging approach (Figure 3C). Consistent with our RIP findings, the TERRA-MDA5 PLA in U2OS cells yielded a significantly higher number of interaction foci than the negative-control probe (Figure 3D-F). Notably, these macromolecular complexes were detected both within the nucleus, where TERRA is natively synthesized and retained, and in the cytoplasm, where the RLR signaling machinery resides (Figure 3E). To bridge these *in vitro* findings with our *in vivo* model, we isolated primary cells directly from RAS zebrafish brains and performed the same RNA-protein PLA assay. Consistently, this tissue-specific analysis recapitulated our cell line data, demonstrating an enrichment of TERRA-MDA5 interaction foci compared to negative-control setups (Figure 3G-I). Because MDA5 strictly recognizes structured or double-stranded RNA (dsRNA) ligands, these binding dynamics imply that a fraction of cytoplasmic TERRA exists in a duplex conformation. To verify this structural requirement, we performed PLA coupling TERRA probes with the J2 antibody, which specifically recognizes dsRNA epitopes^48^. This analysis revealed an enrichment of TERRA-J2 interaction foci relative to negative-control probes (Figure 3J-L), confirming that TERRA forms double-stranded structures compatible with MDA5-mediated surveillance. Given the T and G rich sequence of TERRA, self-pairing of TERRA transcript is unlikely to happen unless it takes place at the level of the subtelomeric regions, where the sequences may be enriched with a more variable nucleotide sequence. Interestingly, it is known that TERRA can interact with TERC (the RNA component of the telomerase holoenzyme^33^), at the level of the template domain. To verify if TERRA-TERC interaction could explain, at least in part, the sensing of double-stranded TERRA by MDA5 we investigated whether TERC too could interact with MDA5 and employed the PLA technique to identify TERC-MDA5 interactions. We found a significantly higher number of TERC-MDA5 foci in ALT+ cells compared with the negative control (Figure 4M-O), suggesting that double-stranded RNA molecules generated by the base-pairing of TERRA and TERC can be sensed by MDA5. This result suggests a possible mechanism linking MDA5 receptors, TERRA and TERC in ALT+ tumors.

**Figure 3:**
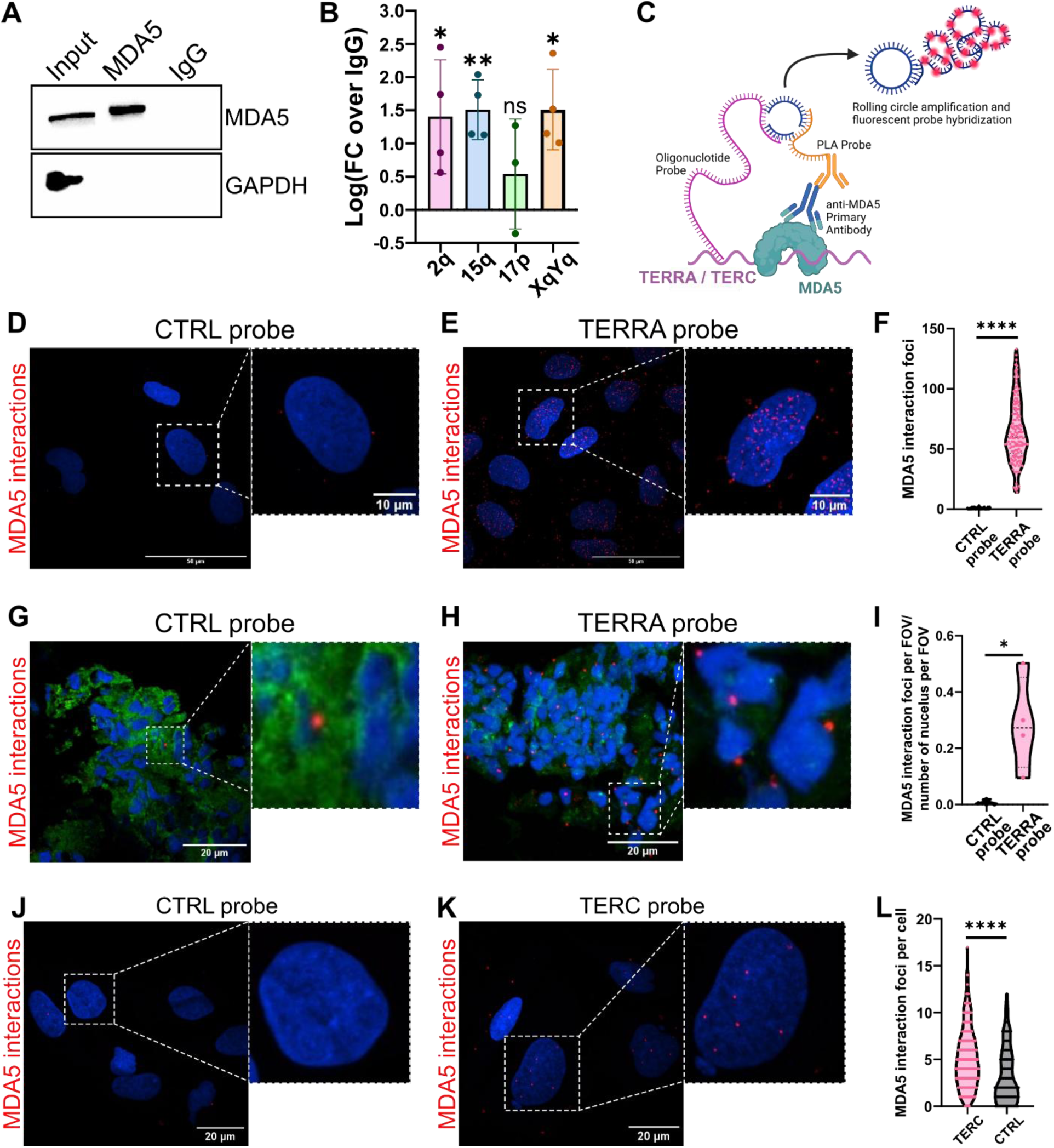
Increased TERRA engages the RLR pathway to induce inflammation. (A) Western blot indicating the successful enrichment of MDA5 in MDA5-pull own sample. (B) Quantification of TERRA coming from specific telomeres (2q, 15q, 17p and XqYq) immunoprecipitated with MDA5, indicated as fold-change enrichment over IgG negative control, in logarithmic scale (Statistics: One sample t-test, n=>3,**p<0.01; *p<0.05; ^ns^p>0.05). (C) Schematic representation of PLA technology used to detect TERRA/TERC interaction with MDA5 (Created with Biorender.com) (D-E) Representative images of PLA performed in U2OS cells with MDA5 antibody and CTRL (D) or TERRA (E) probe. TERRA interaction foci are in red, DAPI is in blue. Images are processed with Fiji. Scale bar: 50 µm and 10 µm. (F) Relative quantification of the MDA5 interaction foci in U2OS (Statistics: Mann Whitney test; CTRL probe n=99, TERRA probe n=141; ****p<0.0001). (G-H) Representative images of PLA performed in Zic:RAS brain sections with MDA5 antibody and CTRL (G) or TERRA (H) probe. TERRA interaction foci are in red, DAPI is in blue, RAS-GFP is in green. Scale bar: 20 µm. (I) Relative quantification of the MDA5 interaction foci in Zic:RAS brain slices (Statistics: Unpaired t-test; n=4; *p<0.05). (J-K) Representative images of PLA performed in U2OS cells slices with MDA5 antibody and CTRL (J) or TERC (K) probe. TERC interaction foci are in red, DAPI is in blue. Scale bar: 20 µm. (L) Relative quantification of the MDA5 interaction foci in U2OS cells (Statistics: Mann Whitney test; CTRL probe n=105, TERRA probe n=214; ****p<0.0001).

**Figure 4:**
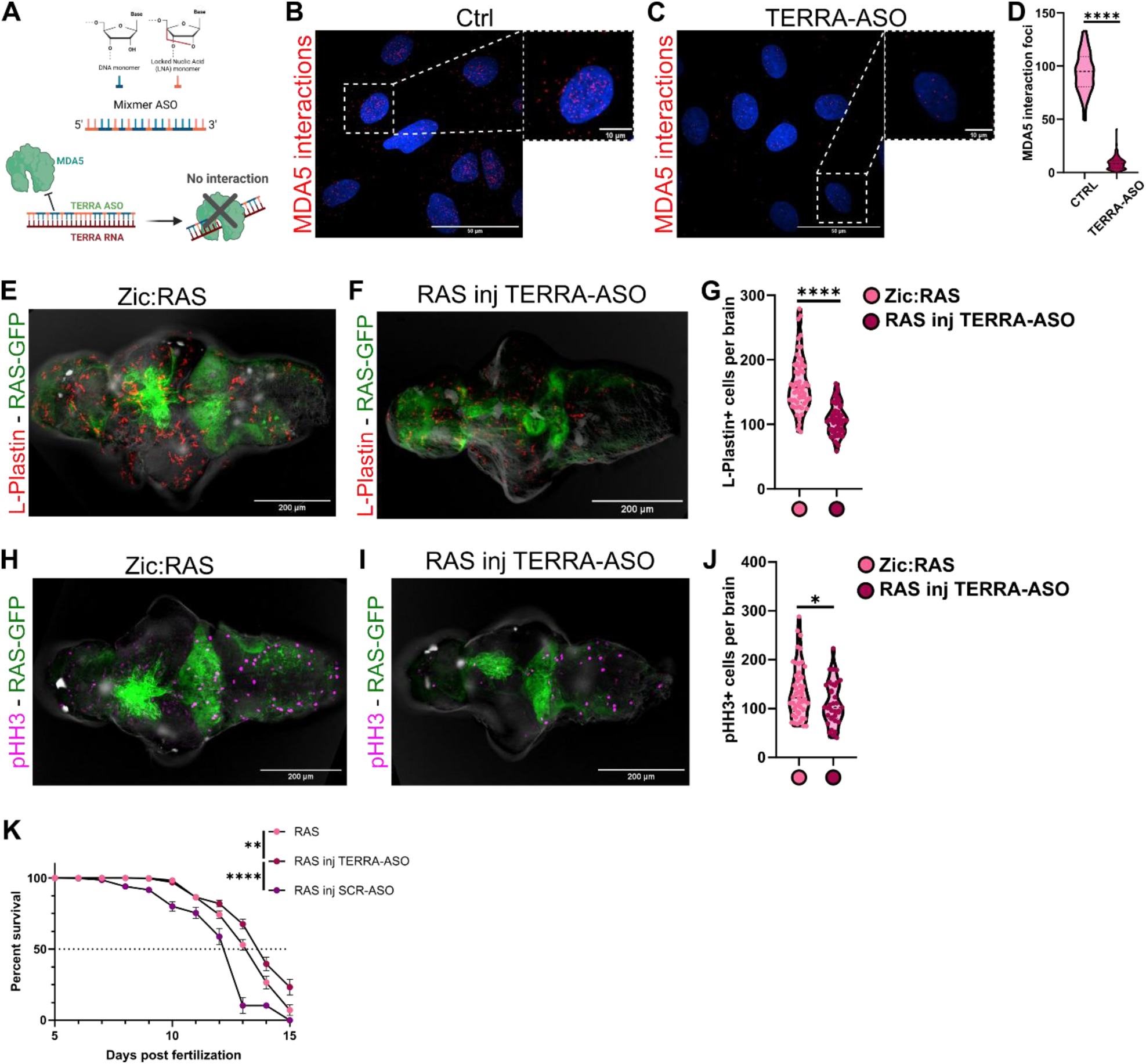
Antisense TERRA reduces inflammation, tumor cell proliferation and prolong survival of tumor bearing larvae. (A) Schematic representation of the TERRA targeting mixmer ASO structure and mechanism of action used in the study. (B-C) Representative images of PLA performed in U2OS cells transfected or not with TERRA-ASO using MDA5 antibody and TERRA probe. TERRA interaction foci are in red, DAPI is in blue. Images are processed with Fiji. Scale bar: 50 µm and 10 µm. (D) Relative quantification of the MDA5 interaction foci in U2OS (Statistics: Mann Whitney test; CTRL n=41, TERRA probe n=108; ****p<0.0001). (E-F) Representative WM-IF images of 5 dpf larval brains of Zic:RAS (E) and RAS inj ASO (F), showing L-plastin staining (red) and RAS-GFP expression (green). Scale bar: 200 μm. (G) Quantification of the number of L-plastin+ cells per brain (Statistics: Kruskal-Wallis test with Dunn’s multiple comparisons test, Zic:RAS n=55; RAS inj ASO n=59; ****p<=0.0001). (H-I) Representative WM-IF images of 5 dpf larval brains of Zic:RAS (H) and RAS inj TERRA-ASO (I), showing pHH3 staining (magenta) and RAS-GFP expression (green). Scale bar: 200 μm. (J) Quantification of the number of pHH3 cells per brain (Statistics: Mann Whitney test, Zic:RAS n=46; RAS inj TERRA-ASO n=34; *p<=0.05). (K) Survival curve of Zic:RAS fish not injected, injected with TERRA-ASO or with SCR-ASO up to 15 dpf (Statistics: Log-rank (Mantel-Cox) test, Zic:RAS n=135; at 5 dpf RAS inj TERRA-ASO n=117; RAS inj SCR-ASO n=60; **p<=0.001; ****p<=0.0001).

### 4. Antisense TERRA reduces inflammation, tumor cell proliferation and prolong survival of tumor bearing larvae

To confirm that TERRA drives MDA5-mediated sterile inflammatory response in ALT+ tumors, we employed an antisense oligonucleotide (ASO) steric blocking strategy. Because TERRA forms highly structured configurations recognized by MDA5, we used a mixmer ASO targeting the repetitive 3’-UUAGGG-5’ tract common to the entire TERRA population (Figure 4A). Unlike gapmer ASOs that trigger RNase H-mediated transcript degradation, which showed no reduction in total TERRA levels in the Zic:RAS model (Supplementary Fig. 4A-B), the mixmer design functions through steric hindrance, masking the RNA to disrupt macromolecular interactions without altering steady-state transcript abundance^60–64^. To confirm the efficacy of the steric blockade, we transfected U2OS cells with the TERRA-ASO and assessed TERRA-MDA5 interaction via TERRA-protein PLA analysis. The number of TERRA-MDA5 interaction foci present after ASO transfection was significantly lower than in the scramble-treated controls (Fig. 4B-D), confirming that the mixmer TERRA-ASO effectively shields TERRA from MDA5. Next, we translated these findings *in vivo* to assess the functional consequences of disrupting the TERRA-MDA5 axis during zebrafish brain tumorigenesis. We microinjected the TERRA-ASO into Zic:RAS embryos at the single-cell stage and evaluated leukocyte infiltration and mitotic activity in the brains of 5 dpf larvae. Larvae receiving TERRA-ASO exhibited a robust reduction in immune cell infiltration (Figure 4E-G) alongside a significant decrease in pHH3+ mitotic cells (Figure 4H-J). At the same time, the injection of TERRA-ASO in healthy fish (supplementary Fig. 4C), as well as the injection of non-targeting SCR-ASO in Zic:RAS embryos (supplementary Fig. 4D), did not alter the number of L-plastin+ cells in the brain confirming a tumor-specific function for TERRA and its targeting ASO. These anti-inflammatory and anti-proliferative effects closely mirrored the phenotypes obtained in MDA5 and MAVS crispants, further substantiating that TERRA acts as an upstream trigger of tumor-associated inflammation through MDA5. Finally, to decipher whether this TERRA-driven immune microenvironment exerts a pro- or anti-tumorigenic effect on overall disease progression, we monitored the survival of RAS juveniles up to 15 dpf. Survival trajectories were compared among untreated RAS fish, those injected with a non-targeting scramble control (SCR-ASO), and those injected with the TERRA-ASO. Notably, altering TERRA-MDA5 interaction conferred a survival advantage, with TERRA-ASO-injected fish exhibiting significantly higher survival rates than both baseline controls and SCR-ASO cohorts (Figure 4K). Taken together, these survival dynamics indicate that the downstream RLR inflammatory cascade behaves in a pro-tumorigenic manner, and that disrupting this specialized RNA-sensing loop effectively reduces tumor progression.

### 5. TERRA ASO treatment affects ALT+ cancer cells in a long-lasting way

Most of the analyses conducted so far relate to 5 dpf animals, in which the tumor is in the early stages of development and the immune system is not completely developed. For this reason, to gain more insight about the role of TERRA ASO in shaping the TME and improving the survival of the fish developing brain tumors we conducted single-cell RNA-seq to compare the transcriptome of the different cells in adult tumors from Zic:RAS or Zic:RAS injected with TERRA ASO (named RAS inj TERRA ASO). A total of 49729 cells were captured from 7 biological replicates of two models (4 *RAS* – 21789 cells, 3 RAS inj TERRA ASO – 27940 cells), comprising both cancer and TME cells (supplementary Fig. 5A, Supplementary table 3). Unsupervised clustering following quality control (see M&M) identified 31 clusters with distinct transcriptional profiles (supplementary 5B, table 4). During cell type annotation, we defined clusters expressing the transgene *(eGFP-HRASV12G,* supplementary Fig*. 5C-D*) as cancer cells and all the other clusters as different cell types of the tumor microenvironment and the surrounding brain. We found 2 clusters of cancer cells that differ because one of them is strongly characterized by the expression of *prss35* and *gpr37L1A*, two markers of Bergman glia^65^, and we interpret this gene expression as a marker of their location: the cancer cells in this group belong to cerebellar tumors. Therefore, we named them as cancer cell 1 (cancer cells from tumors in the forebrain) and cancer cell 2 (cancer cells from the hindbrain region). We merged all the other clusters that expressed the same genes to obtain a final list of 18 cell types (see M&M, Fig. 5A, B, supplementary table 4). The cell types identified in our analysis were: cancer cells, macrophpages/microglia cells, NK/T cells, endothelial cell, pericytes, astrocytes, oligodendrocytes, oligodendrocyte progenitor cells, radial glia cells, quiescent radial glia cells, fibroblast, lymphatic endothelial cells, B cells, T-helper 2 cells, neurons, dendritic cells and neural progenitor proliferative cells. The different cell types expressed specific markers, reported in M&M. To identify the differences in cancer cells between Zic:RAS and RAS inj TERRA-ASO we subclustered them (Figure 5C and Supplementary table 5), obtaining 8 cancer cell subclusters. By comparing the relative abundance of cells from Zic:RAS and RAS inj TERRA-ASO distributed in the 8 clusters, we found that cluster 2 is significantly overrepresented in Zic:RAS, whereas clusters 4 is significantly larger in RAS inj TERRA-A*SO* (Supplementary 5E and table 6). To facilitate the subsequent analysis, from now on, cluster 2 will be named C2-RAS, and cluster 4 will be referred to as C4-ASO. We used these 2 clusters to capture the most significant differences induced by TERRA ASO treatment and carried out differential expression (DE) analysis and gene set enrichment analysis (GSEA) to identify them. From these analyses it emerged that 296 genes were significantly upregulated in C4-ASO compared to C2-RAS, whereas 136 genes were downregulated (Figure 5D). In particular in C4-ASO, genes related to synaptic trafficking and differentiation (*robo1, dlgap2a, shank1, shank3b, nlgn1, nlgn2a, nlgn3a, ntrk3a, grin2b, grik5, grid2, grm8a, ntm, pcdh1a, cadm1a, camkk1a, l1cama, kcnd2, kcnn4, hcn5, cacna1i, cacnb3a, sema3e, plxna1a, plxna4, nrp2a, slitrk3a, auts2a, dpyda.1, ntn4, rtn4r, rtn4rl1a* – in purple in Fig. 5D) were overexpressed. In contrast, C2-RAS is characterized by the expression of genes related to epithelial to mesenchymal transition (EMT), extracellular matrix remodeling (*anxa5b, anxa1a, anxa2a, vim, tagln, tuba1a, tuba1c, tubb4b, ahnak, calm2b, cav1, cavin1b*, *mmp9, fn1a, col12a1a, col15a1a, col1a2, mfap5, plpp3, prss23, adam8a* – in yellow in Fig. 5D), and stemness (*sox4a, id1* – in yellow in Fig. 5D), suggesting a more aggressive and invasive phenotype of this cluster of cancer cells. A large group of genes downregulated in C4-ASO belongs to the cell’s translational machinery, such as ribosomal proteins (*rpl11, rpl19, rpl5b, rpl7, rps3a, rps4x, rplp2l, rps27l, rps12, rpl15, rps6, rpl23a, eef1da, tpt1, npm1a, snu13b, tomm5* – in green in Fig. 5D) and proteasome subunits (*psme1, psmb4, psmb3, psmb2* – in green in Fig. 5D). A drop in protein synthesis and turnover correlates with a reduction in biomass production, which may be linked to reduced proliferation^66^. In line with this, genes involved in apoptosis, in the stress response program, and consequently in growth arrest (*pdcd4b, trib3, rasa2, rassf5, foxn3*, *fosb, nr4a3, btg1, btg2, crema, csrnp1b, csrnp3, dusp5, ppp1r15a, gab2* - in pink in Fig.5D) were upregulated in C4-ASO. Taken together, this differential gene expression analysis suggests that C4-ASO cancer cells, compared to C2-RAS cancer cells, are less proliferative and less invasive, less stem-like, and are presumably less aggressive. Moreover, in C4-ASO we also found downregulated inflammation-related genes (*isg15, ifi46, cxcl14, cxcl18b, lgals2a, mt2* – in orange in Fig. 5D), further supporting the idea that TERRA possesses immunostimulatory properties and that targeting these transcripts in tumors may have a broad spectrum of effects not only related to telomere maintenance. Limiting this analysis to two cancer cell populations, each one representative of one model, allowed us to highlight the most relevant differences between Zic:RAS and RAS inj TERRA-ASO cancer cells. However, given the decreased proliferation that we detected in RAS inj TERRA-ASO tumors, we extended the cell cycle status analysis to the entire population of cancer cell. In line with the DE analysis, we found that C4-ASO cancer cells were cycling less compared to those of the C2-RAS cluster, with a percentage of cells in G1 of 70.2% compared to 45.3% in C4-ASO (supplementary Fig. 5E). This proportion is also reflected in the models, indeed by clustering cancer cells according to the models, upon ASO treatment the percentage of cells in G1 goes from 38.9% to 57.4% (Figure 5E), indicating that in RAS inj TERRA-ASO cancer cells are replicating less. In support of these results, GSEA of Hallmark gene sets (Figure 5F, Supplementary table 7) and Reactome pathways^66^ analysis (Figure 5G, Supplementary table 8) highlighted a similar trend of reduced protein synthesis and cell proliferation, along with increased neuronal communication in RAS inj TERRA-ASO. Of note, Reactome GSEA also highlighted a reduction in the “viral infection pathway” in C4-ASO, whereas it was increased in C2-RAS, suggesting a possible shutdown or downregulation of genes downstream of MDA5 activation.

**Figure 5:**
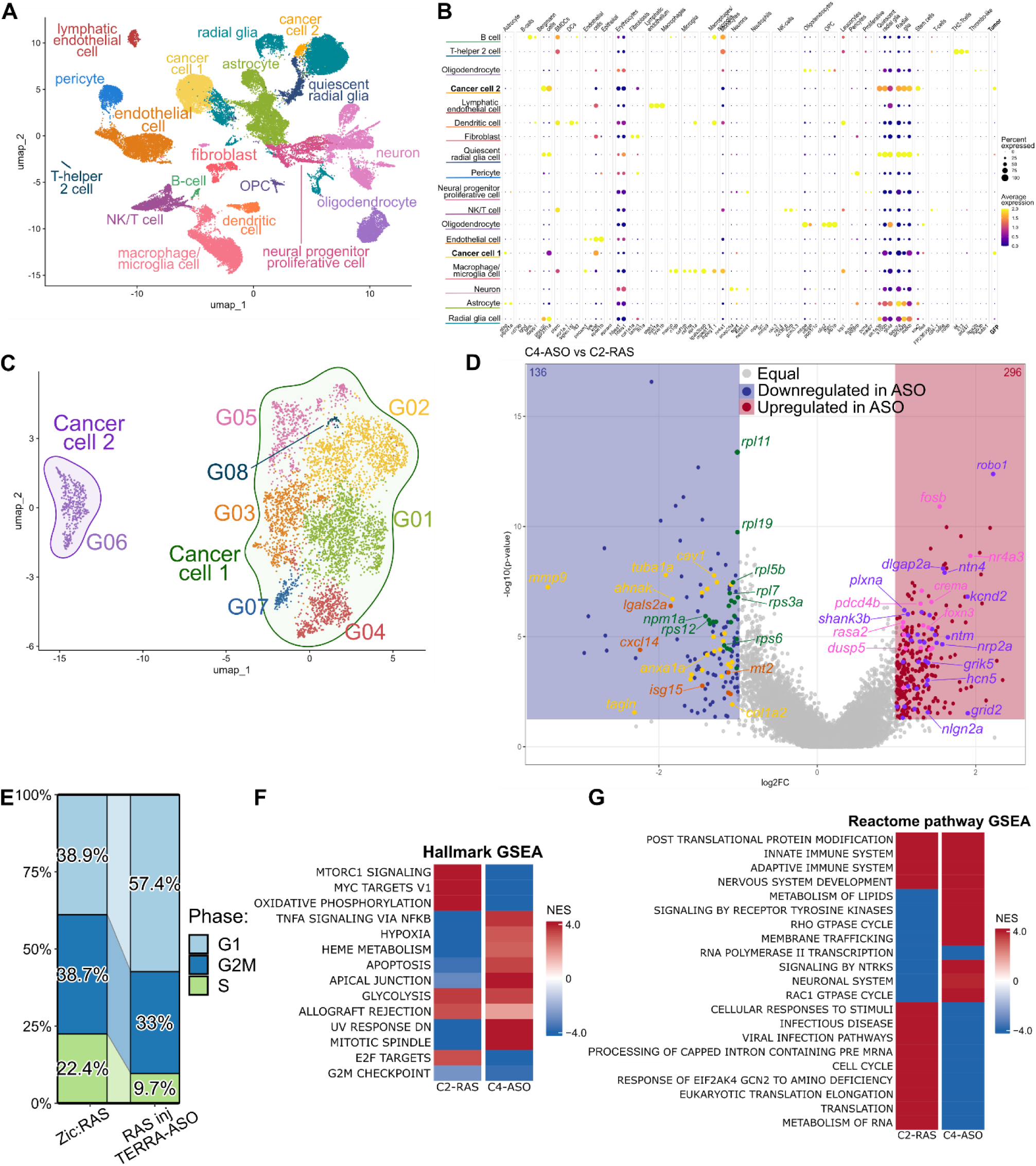
TERRA ASO treatment affects ALT+ cancer cells in a long-lasting way. (A) Uniform manifold approximation projection (MAP) of 49729 aggregated cells coming from the two models, showing the different cell types composing glioblastoma in zebrafish, highlighting tumor clusters (cancer cell 1, cancer cell 2), neural lineages (radial glia, astrocytes, neurons, OPCs, oligodendrocytes), vascular elements, and immune populations (microglia/macrophages, T/NK cells, B cells).(B) Dot plot showing the expression of marker genes (x-axis) used for the manual annotation of the major cell types (y-axis) found in the dataset. Dot size indicates the percentage of cells of a given cell type expressing the gene and the color indicating the average expression ( high expression in yellow and low expression in blue). (C) UMAP projection of *de novo* clustering of the cancer cells from the two models showing the 8 clusters obtained from unsupervised clustering grouped into two main cancer cell states (Cancer cell 1 highlighted in purple and Cancer cell 2 highlighted in green). (D) Volcano plot showing the differentially expressed genes comparing the C4-ASO subcluster against C2-RAS, highlighting in blue 136 downregulated and in red 296 upregulated genes. In green genes involved in protein synthesis, in yellow EMT-related genes, in orange ISG, in pink genes correlated with cell cycle arrest and in purple genes for trafficking and differentiation – see text for more details. (E) Stacked bar plot comparing the percentages of cancer cells in the G1, G2M or S cell cycle phase in the Zic:RAS and Zic:RAS ASO models. (F-G) Heatmaps displaying Normalized Enrichment Scores (NES) from Gene set enrichment analysis (GSEA) comparing the C2-RAS cluster and C4-ASO clusters using (F) Hallmark gene sets and (G) Reactome pathways. The color scale represents activation (red, NES > 0) or suppression (blue, NES < 0) of key biological processes, emphasizing the downregulation of proliferation-related programs (E2F targets, G2M checkpoint, cell cycle) alongside modulation of innate/adaptive immune pathways and metabolic states in C4-ASO.

### 6. Immune cell populations are affected by ASO targeting TERRA in ALT+ brain tumors

Given our data showing a change in microglial and macrophage population in the brains of 5 dpf Zic:RAS larvae following MDA5/MAVS KOs or after TERRA ASO injection we were intrigued to know what changes in the immune cells infiltrating Zic:RAS brain tumors were induced by ASO targeting TERRA. We took advantage of the single-cell data to analyze the immune cell populations in Zic:RAS and RAS inj TERRA-ASO tumors. De novo clustering of immune cells populations identified 7 cluster corresponding to the major cell type: T-helper 2 cells (expressing *il4*, *il13* and *cd247*), neutrophils (expressing *mpx*, *lyz* and *mmp9*), NK cells (expressing *nkl.3*, *nkl.4*, *ccl38.6*, *il2rb* and *gzm3.3*), T cell (expressing *FP236356.1*, *cd4-1*, *cd8a* and *cd8b*), B-cells (expressing *cd79b*, *ighd*, *blnk* and *dapp1*), DCs (expressing *xcr1a.1*, *singlec15l* and *flt3*) and macrophages/microglia cells (expressing *marco*, *c1qb*, *csf1rb*, *csf1ra*, *spi1a*, *lgals3bpb*, *mpeg1.2* and *mpeg1.1*) (Figure 6A-B, Supplementary table 9-10). Subclustering allowed us to compare the proportion of each cell type between the two models (Figure 6C). The two cell populations that changed the most upon ASO injection were the macrophages/microglia cells (reduced after ASO injection), and T-cells (increased in ASO injected brain tumors). Since macrophages/microglia (from here referred as macs/mics) are the most abundant immune cell population in zebrafish brain we focused our attention on these cells and performed a differential expression analysis comparing the mac/mic population present in RAS inj TERRA-ASO with the same cell population in Zic:RAS (Figure 6D, supplementary table 11). From this analysis emerged that 350 genes were upregulated in macs/mics in RAS inj TERA-ASO, while 89 were downregulated compared to non-injected Zic:RAS. We observed a significant reduction in the expression of ISGs and inflammatory signals (*isg15, irf7, cxcl11.5, cxcl11.6, cxcl8a, tnfb, nfkbiaa*), together with TAM markers (*spp1, serpine1, grn1, sqstm1, anxa3b, psap*)^65,66^ after ASO treatment. At the same time, there was in increase in transcripts associated with cell motility (*myo9ab, sptbn5, marcksl1a, mtss1la, arhgap20b, arhgap29a, raph1b, carmil3, cyth3a*), suggesting a migratory or morphologically dynamic cell state rather than a purely inflammatory one, as well as genes involved in surveillance and homeostasis (*foxo1b, foxo3b, bach2b*)^67^. Thus, what emerged from this analysis is that in RAS inj TERRA-ASO tumors, the macs/mics lose their inflammatory and TAM signatures, while acquiring a more homeostatic status. These data are in line with the idea that the ASO treatment impairs the activation of an immune pathway in cancer cells, which function as a macrophage-recruiting or re-educating signal. Overall, the ASO treatment appears to disengage the macrophage population from the tumor-induced inflammatory program and revert them toward a more homeostatic, surveillance-like, and motile state. Together with our findings in cancer cells and survival analysis, these data suggest that TERRA ASO treatment blunts TERRA-mediated immune responses in cancer cells. This reduces the inflammatory signals within the TME, and could favor a more functional, anti-tumoral immune response that ultimately yield a favorable therapeutic outcome.

**Figure 6:**
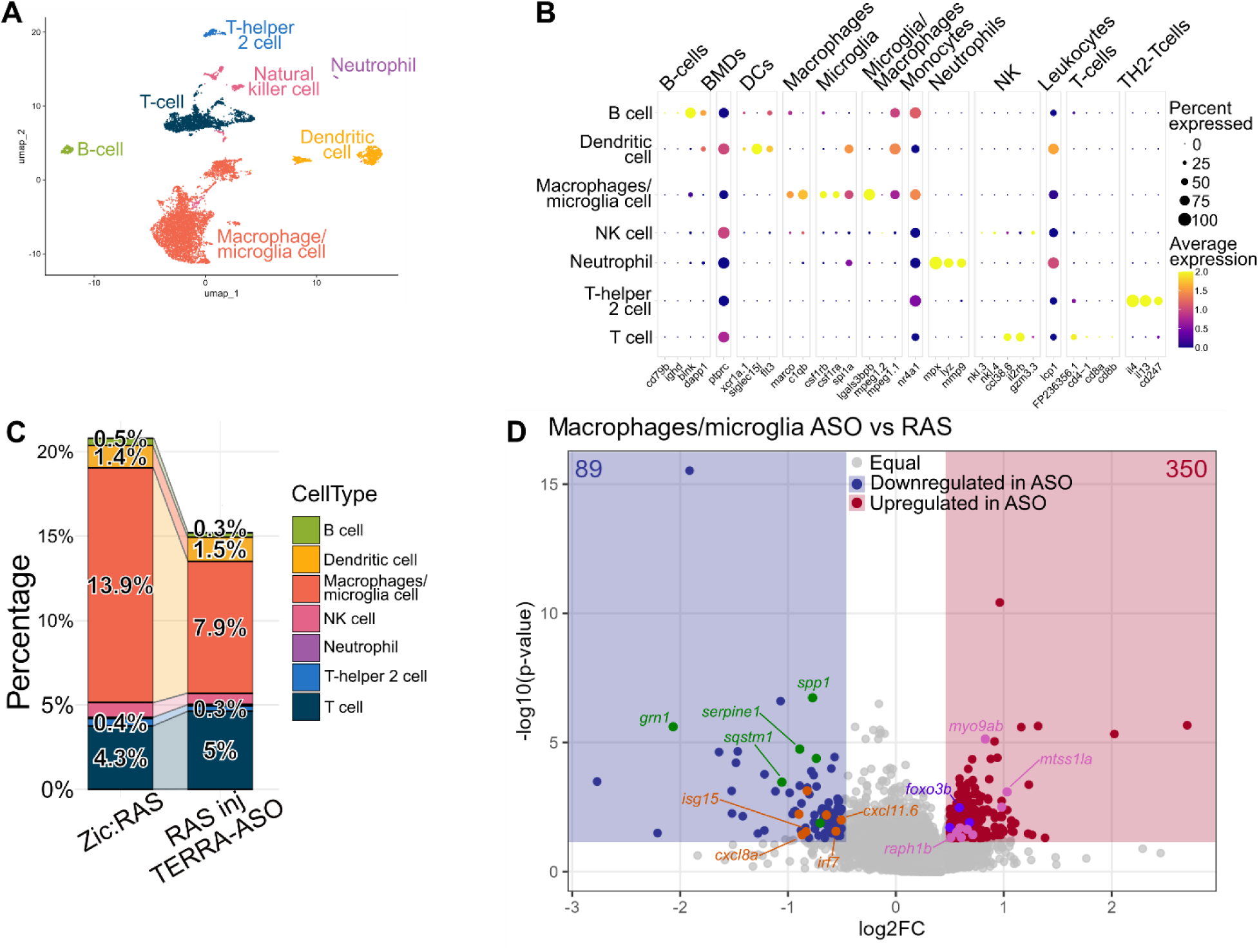
Immune cell populations are affected by ASO targeting TERRA in ALT+ brain tumors. (A) UMAP projection of de novo clustering of 8686 immune cells presents in the two models showing the major cell types found in the dataset. (B) Dotplot showing the genes used for manual annotation of the immune cell types found in the dataset. Dot size indicates the percentage of cells expressing the gene while the color indicates the average expression (high expression in yellow and low expression in blue) (C) Stacked barplot showing the percentages of immune cells in each model (normalized to the total number of cells in that model) comparing the Zic:RAS and RAS inj TERRA-ASO. (D) Volcano plot displaying differentially expressed genes in macrophages/microglia (RAS inj TERRA-ASO vs Zic:RAS), highlighting in blue 89 downregulated and in red 350 upregulated genes. In green genes related to TAM phenotype, in orange ISGs, in pink cell-motility associated genes, in purple surveillance genes – see text for more details

## Discussion

The acquisition of replicative immortality through telomerase reactivation or Alternative Lengthening of Telomeres (ALT) is a fundamental hallmark of cancer. Beyond supporting sustained proliferation, telomere maintenance mechanisms influence tumor biology by impacting genome stability and cellular stress responses. ALT is characterized by telomere dysfunction, elevated telomeric recombination, and increased expression of the long non-coding RNA TERRA, raising the possibility that ALT-associated changes directly influence the tumor microenvironment (TME). Using a zebrafish model of ALT+ brain tumors, we identify TERRA as a key mediator linking ALT-associated telomere biology to innate immune activation and tumor progression. We observed that TERRA upregulation accompanies the early stages of tumorigenesis and persists in fully developed tumors, co-occurring with the upregulation of inflammatory cytokines, NF-κB signaling and type I ISG expression, with a substantial infiltration of L-plastin-positive immune cells of the tumours. These findings suggest that inflammatory signaling is established early during tumorigenesis and persists in fully developed tumors. Although the coexistence of TERRA accumulation and inflammation does not, by itself, establish a causal relationship, our genetic and antisense approaches to modulate this pathway support a functional contribution of TERRA-dependent RNA sensing to this response. Recent work has implicated TERRA in innate immune activation through ZBP1 binding^35^. In our zebrafish model, we found evidence supporting the involvement of TERRA-MDA5 interaction in tumor development. Mechanistically, our genetic and biochemical experiments demonstrate that TERRA engages the double-stranded RNA sensor MDA5 and its downstream effector MAVS. Overexpression of MDA5 in tumor cells enhanced immune infiltration and tumor proliferation, possibly due to a prolonged and dysregulated activation of type I interferon responses. On the contrary, CRISPR/Cas9-mediated disruption of *mda5* or *mavs* attenuated both immune infiltration and cell proliferation. In contrast, RIG-I failed to reproduce these effects, highlighting sensor specificity within the tumor context. Furthermore, RNA-RIP and PLA assays in U2OS cells and primary zebrafish tumor cells confirmed a physical association between TERRA and MDA5, with J2 antibody recognition indicating that a fraction of TERRA adopts a double-stranded immunogenic conformation, potentially through secondary structures or interactions with other cellular transcripts such as TERC. However, proximity assays do not establish direct molecular binding or demonstrate that TERRA-TERC duplexes are the ligands responsible for MDA5 activation. Defining the sequence, structure, subcellular origin, and functional properties of the RNA species recognized by MDA5 in this context will therefore be an important direction for future investigation. A central finding of this study is that interfering with TERRA attenuates the inflammatory response and reduces tumor-cell proliferation. Targeting this signaling axis using a steric-blocking mixmer ASO against TERRA’s telomeric repeats effectively disrupted TERRA-MDA5 binding, reduced leukocyte infiltration, decreased mitosis in tumors, and improved larval survival without affecting healthy fish. The phenotypic similarities between ASO-injected larvae and MDA5- or MAVS-deficient larvae, supports the interpretation that TERRA contributes to the inflammatory phenotype. Furthermore, the improved survival of ASO-injected tumor-bearing larvae suggests that the inflammatory response associated with this pathway has adverse consequences for the host. Nevertheless, because the ASO is administered at the single-cell stage and may influence TERRA interactions in multiple cellular compartments, additional experiments will be needed to establish the relative contributions of cancer-cell-intrinsic and immune-cell-intrinsic effects. Single-cell transcriptomic analysis further revealed that early embryonic ASO treatment drives persistent remodeling in adult tumors: cancer cells shifted toward a less proliferative, non-stem-like state with reduced epithelial-to-mesenchymal transition and extracellular matrix remodeling programs. These changes were accompanied by a greater proportion of cells in G1 and reduced cycling cells, consistent with the lower mitotic activity observed in larvae. Tumor-associated macrophages and microglia in ASO-injected fish exhibited diminished inflammatory and interferon-responsive signatures alongside a relative increase in T-cell representation. These changes suggest that TERRA-associated signaling influences both the recruitment or persistence of myeloid cells and their transcriptional state. In particular, the reduction in inflammatory and tumor-associated macrophage signatures is consistent with a model in which tumor-cell RNA sensing contributes to the establishment or maintenance of a pro-tumoral myeloid environment. Further studies combining lineage-specific manipulation, spatial analysis and functional assessment of immune populations will be required to determine how TERRA-dependent signaling coordinates communication between tumor cells and the immune microenvironment. The persistence of these effects in adult tumors following early TERRA-ASO treatment suggests that interrupting RNA sensing during embryogenesis alters the long-term trajectory of tumor growth and microenvironmental organization. Because adult tumors were compared to untreated controls and analyzed via single-cell transcriptomic snapshots, it remains unclear whether this outcome reflects a durable direct effect of TERRA inhibition or a secondary consequence of reduced early tumor burden. Temporally controlled interventions and longitudinal studies are needed to determine whether TERRA signaling is continuously required throughout tumor progression or acts primarily during an early developmental window. In this model, the proposed sequence of events involves increased TERRA expression, engagement of MDA5 signaling, induction of inflammatory programs, remodeling of the immune microenvironment and enhanced tumor-cell proliferation. The functional effects of MDA5/MAVS disruption and TERRA-targeting ASOs support several steps in this model, but the precise molecular intermediates connecting RNA sensing to immune recruitment and cancer-cell proliferation remain to be determined. In conclusion, our study identifies TERRA-associated MDA5 signaling as a contributor to inflammation and tumor progression in a zebrafish model of ALT+ brain tumors. By combining genetic manipulation, molecular interaction assays, functional TERRA inhibition and single-cell transcriptomics, we show that interference with this pathway is associated with reduced immune infiltration, decreased tumor-cell proliferation, improved larval survival and persistent changes in cancer-cell and myeloid-cell transcriptional states. These findings support a model in which TERRA participates in linking ALT-associated telomere biology to the tumor microenvironment and suggest that targeting TERRA-mediated RNA sensing may offer a strategy to modulate tumor-associated inflammation in ALT-positive cancers.

## Acknowledgements

The authors would like to thank Veronica Bergo and Stylianos Lefkopoulos (previous members of the Trompouki’s lab) for generating the transgenic construct used for the production of the UAS:hsaMDA5 zebrafish line. We thank Dr. Chien’s laboratory (University of Utah, USA) for developing and sharing the Tol2kit vectors. Plasmids used in this study were obtained from the Tol2kit website. We thank Emilio Cusanelli (CIBIO Department, University of Trento) for advice in studying TERRA in zebrafish. We thank members of the Core Facilities (CIBIO Department, University of Trento) for their participation in quality control, animal care and sequencing steps, and all members of the Mione laboratory for their help and insightful discussions.

## Author Contributions

**Pecchini:** conceptualization, data curation, formal analysis, supervision, validation, investigation, visualization, methodology, writing–original draft, writing–review and editing. **V. Bontempi:** data curation, software, formal analysis, validation, investigation, methodology, visualization, funding acquisition, writing– original draft, writing–review and editing. **C. Isidori**: investigation, data curation, formal analysis, validation. **S. Sorze:** methodology, visualization. **A. Liuzzo:** methodology, visualization. **Inmaculada Martínez Olmo:** methodology, visualization. **Maria Luisa Cayuela:** supervision**. N. Abdelfattah:** software, data curation, formal analysis. **T. Tebaldi:** software, data curation, supervision, writing–review and edit. **K. Yun:** resources, data curation, supervision, writing–review and editing. **E. Trompouki:** resources, supervision, writing–review and editing. **M.C. Mione:** conceptualization, resources, data curation, formal analysis, supervision, funding acquisition, validation, investigation, visualization, methodology, writing–original draft, project administration, writing–review and editing.

## Language editing

Large language models, including ChatGPT (OpenAI), were used to assist in improving the clarity and fluency of English in the manuscript text. These tools were not used for data analysis, interpretation, or generation of scientific content.

## List of abbreviations

ALT: Alternative Lengthening of Telomeres
APB: ALT-associated PML body
ASO: Antisense Oligonucleotide
CCF: Cytoplasmic Chromatin Fragment
DC: Dendritic Cells
DPF: Days post fertilization
GBM: Glioblastoma
HGG: High Grade Glioma
IF: Immunofluorescence
ISG: Interferon Stimulated Gene
lncRNA: Long Non-Coding RNA
PLA: Proximity Ligation Assay
PRR: Pattern-Recognition Receptors
RIP: RNA Immunoprecipitation
TAM: Tumor Associated Macrophages
TERRA: Telomeric Repeat containing RNA
TME: tumor microenvironment
TMM: Telomere Maintenance Mechanism
WM: Whole-Mount

## References

1. O’Sullivan, R. J. & Karlseder, J. Telomeres: protecting chromosomes against genome instability. Nat. Rev. Mol. Cell Biol. 11, 171–181 (2010).

2. Sfeir, A. J., Chai, W., Shay, J. W. & Wright, W. E. Telomere-End Processing: the Terminal Nucleotidesof Human Chromosomes. Mol. Cell 18, 131–138 (2005).

3. Makovets, S., Herskowitz, I. & Blackburn, E. H. Anatomy and Dynamics of DNA Replication Fork Movement in Yeast Telomeric Regions. Mol. Cell. Biol. 24, 4019–4031 (2004).

4. Verdun, R. E. & Karlseder, J. The DNA Damage Machinery and Homologous Recombination Pathway Act Consecutively to Protect Human Telomeres. Cell 127, 709–720 (2006).

5. Opresko, P. L., Fan, J., Danzy, S., Wilson, D. M. & Bohr, V. A. Oxidative damage in telomeric DNA disrupts recognition by TRF1 and TRF2. Nucleic Acids Res. 33, 1230–1239 (2005).

6. Hoffelder, D. R. et al. Resolution of anaphase bridges in cancer cells. Chromosoma 112, 389–397 (2004).

7. Maciejowski, J., Li, Y., Bosco, N., Campbell, P. J. & de Lange, T. Chromothripsis and kataegis induced by telomere crisis. Cell 163, 1641–1654 (2015).

8. Chen, Y.-A. et al. Extrachromosomal telomere repeat DNA is linked to ALT development via cGAS-STING DNA sensing pathway. Nat. Struct. Mol. Biol. 24, 1124–1131 (2017).

9. Hanahan, D. & Weinberg, R. A. Hallmarks of Cancer: The Next Generation. Cell 144, 646–674 (2011).

10. Tornesello, M. L. et al. Reactivation of telomerase reverse transcriptase expression in cancer: the role of TERT promoter mutations. Front. Cell Dev. Biol. 11, (2023).

11. Dilley, R. L. & Greenberg, R. A. ALTernative Telomere Maintenance and Cancer. Trends Cancer 1, 145–156 (2015).

12. Muoio, D. & Fouquerel, E. The alternative lengthening of telomeres pathway through a DNA repair lens: mechanism and therapeutic opportunities. NAR Cancer 7, zcaf056 (2025).

13. Cesare, A. J. et al. Spontaneous occurrence of telomeric DNA damage response in the absence of chromosome fusions. Nat. Struct. Mol. Biol. 16, 1244–1251 (2009).

14. Bryan, T. M., Englezou, A., Dalla-Pozza, L., Dunham, M. A. & Reddel, R. R. Evidence for an alternative mechanism for maintaining telomere length in human tumors and tumor-derived cell lines. Nat. Med. 3, 1271–1274 (1997).

15. Yeager, T. R. et al. Telomerase-negative Immortalized Human Cells Contain a Novel Type of Promyelocytic Leukemia (PML) Body. https://aacrjournals.org/cancerres/article/59/17/4175/505416/Telomerase-negative-Immortalized-Human-Cells.

16. Azzalin, C. M. TERRA and the alternative lengthening of telomeres: a dangerous affair. FEBS Lett. 599, 157–165 (2025).

17. Episkopou, H. et al. Alternative Lengthening of Telomeres is characterized by reduced compaction of telomeric chromatin. Nucleic Acids Res. 42, 4391–4405 (2014).

18. Arnoult, N., Van Beneden, A. & Decottignies, A. Telomere length regulates TERRA levels through increased trimethylation of telomeric H3K9 and HP1α. Nat. Struct. Mol. Biol. 19, 948–956 (2012).

19. Arora, R. & Azzalin, C. M. Telomere elongation chooses TERRA ALTernatives. RNA Biol. 12, 938–941 (2015).

20. Misino, S., Busch, A., Wagner, C. B., Bento, F. & Luke, B. TERRA increases at short telomeres in yeast survivors and regulates survivor associated senescence (SAS). Nucleic Acids Res. 50, 12829–12843 (2022).

21. Zeinoun, B., Teixeira, M. T. & Barascu, A. TERRA and Telomere Maintenance in the Yeast Saccharomyces cerevisiae. Genes 14, 618 (2023).

22. Manzato, C. et al. TERRA expression is regulated by the telomere-binding proteins POT-1 and POT-2 in Caenorhabditis elegans. Nucleic Acids Res. 51, 10681–10699 (2023).

23. Azzalin, C. M., Reichenbach, P., Khoriauli, L., Giulotto, E. & Lingner, J. Telomeric Repeat–Containing RNA and RNA Surveillance Factors at Mammalian Chromosome Ends. Science 318, 798–801 (2007).

24. Schoeftner, S. & Blasco, M. A. Developmentally regulated transcription of mammalian telomeres by DNA-dependent RNA polymerase II. Nat. Cell Biol. 10, 228–236 (2008).

25. Luke, B. et al. The Rat1p 5′ to 3′ Exonuclease Degrades Telomeric Repeat-Containing RNA and Promotes Telomere Elongation in Saccharomyces cerevisiae. Mol. Cell 32, 465–477 (2008).

26. Diman, A. & Decottignies, A. Genomic origin and nuclear localization of TERRA telomeric repeat-containing RNA: from Darkness to Dawn. FEBS J. 285, 1389–1398 (2018).

27. Feretzaki, M., Renck Nunes, P. & Lingner, J. Expression and differential regulation of human TERRA at several chromosome ends. RNA 25, 1470–1480 (2019).

28. Silva, B., Arora, R., Bione, S. & Azzalin, C. M. TERRA transcription destabilizes telomere integrity to initiate break-induced replication in human ALT cells. Nat. Commun. 12, 3760 (2021).

29. Azzalin, C. M. & Lingner, J. Telomeres: The silence is broken. Cell Cycle 7, 1161–1165 (2008).

30. Xu, M. et al. TERRA-LSD1 phase separation promotes R-loop formation for telomere maintenance in ALT cancer cells. Nat. Commun. 15, 2165 (2024).

31. Dinoi, F. et al. Modulating TERRA G-quadruplexes with ligands: impact on telomeric DNA:RNA hybrids and ALT mechanisms. Nucleic Acids Res. 53, gkaf1300 (2025).

32. Deng, Z., Norseen, J., Wiedmer, A., Riethman, H. & Lieberman, P. M. TERRA RNA Binding to TRF2 Facilitates Heterochromatin Formation and ORC Recruitment at Telomeres. Mol. Cell 35, 403–413 (2009).

33. Bettin, N. et al. TERRA transcripts localize at long telomeres to regulate telomerase access to chromosome ends. Sci. Adv. 10, eadk4387.

34. Redon, S., Reichenbach, P. & Lingner, J. The non-coding RNA TERRA is a natural ligand and direct inhibitor of human telomerase. Nucleic Acids Res. 38, 5797–5806 (2010).

35. Nassour, J. et al. Telomere-to-mitochondria signalling by ZBP1 mediates replicative crisis. Nature 614, 767–773 (2023).

36. Traver, D., Herbomel, P., Patton, E. E., Murphey, R. D. & Et., A. The Zebrafish as a Model Organism to Study Development of the Immune System. Adv. Immunol. 10.1016/s0065-2776(03)81007-6 (2003) doi:10.1016/s0065-2776(03)81007-6.

37. Traver, D. Cellular Dissection of Zebrafish Hematopoiesis.

38. Berman, J., Hsu, K. & Look, A. T. Zebrafish as a model organism for blood diseases. Br. J. Haematol. 123, 568–576 (2003).

39. Carneiro, M. C., de Castro, I. P. & Ferreira, M. G. Telomeres in aging and disease: lessons from zebrafish. Dis. Model. Mech. 9, 737–748 (2016).

40. Hason, M. & Bartůněk, P. Zebrafish Models of Cancer—New Insights on Modeling Human Cancer in a Non-Mammalian Vertebrate. Genes 10, 935 (2019).

41. Raby, L., Völkel, P., Le Bourhis, X. & Angrand, P.-O. Genetic Engineering of Zebrafish in Cancer Research. Cancers 12, 2168 (2020).

42. Kobar, K., Collett, K., Prykhozhij, S. V. & Berman, J. N. Zebrafish Cancer Predisposition Models. Front. Cell Dev. Biol. 9, (2021).

43. Mayrhofer, M. et al. A novel brain tumour model in zebrafish reveals the role of YAP activation in MAPK/PI3K induced malignant growth. Dis. Model. Mech. dmm.026500 (2016) doi:10.1242/dmm.026500.

44. Idilli, A. I. et al. Expression of tert Prevents ALT in Zebrafish Brain Tumors. Front. Cell Dev. Biol. 8, 65 (2020).

45. Sung, J.-Y. Alternative Lengthening of Telomeres: A Prognostic Paradox in Cancer. Cells 14, 1613 (2025).

46. Zhao, W. et al. Exploring tumor-associated macrophages in glioblastoma: from diversity to therapy. *Npj Precis*. Oncol. 9, 126 (2025).

47. D’Alessio, A., Proietti, G., Sica, G. & Scicchitano, B. M. Pathological and Molecular Features of Glioblastoma and Its Peritumoral Tissue. Cancers 11, 469 (2019).

48. Bou-Nader, C. et al. Structural basis of double-stranded RNA recognition by the J2 monoclonal antibody. Nat. Commun. 17, 635 (2025).

49. Jessa, S. et al. K27M in canonical and noncanonical H3 variants occurs in distinct oligodendroglial cell lineages in brain midline gliomas. Nat. Genet. 54, 1865–1880 (2022).

50. Waitkus, M. S., Erman, E. N., Reitman, Z. J. & Ashley, D. M. Mechanisms of telomere maintenance and associated therapeutic vulnerabilities in malignant gliomas. Neuro-Oncol. 26, 1012–1024 (2024).

51. Abdelfattah, N. et al. Single-cell analysis of human glioma and immune cells identifies S100A4 as an immunotherapy target. Nat. Commun. 13, 767 (2022).

52. ZFIN Gene: igf2bp1. https://zfin.org/ZDB-GENE-060929-1258.

53. Bartas, M., et al. Searching for New Z-DNA/Z-RNA Binding Proteins Based on Structural Similarity to Experimentally Validated Zα Domain. Int. J. Mol. Sci. 23, 768 (2022).

54. Rothenburg, S. et al. A PKR-like eukaryotic initiation factor 2α kinase from zebrafish contains Z-DNA binding domains instead of dsRNA binding domains. Proc. Natl. Acad. Sci. U. S. A. 102, 1602–1607 (2005).

55. DRSC - DRSC Integrative Ortholog Prediction Tool. https://www.flyrnai.org/cgi-bin/DRSC_orthologs.pl.

56. Azzalin, C. M. & Lingner, J. Telomere functions grounding on *TERRA firma*. Trends Cell Biol. 25, 29– 36 (2015).

57. Maicher, A., Kastner, L. & Luke, B. Telomeres and disease: Enter TERRA. RNA Biol. 9, 843–849 (2012).

58. Porro, A., Feuerhahn, S., Reichenbach, P. & Lingner, J. Molecular Dissection of Telomeric Repeat-Containing RNA Biogenesis Unveils the Presence of Distinct and Multiple Regulatory Pathways. Mol. Cell. Biol. 30, 4808–4817 (2010).

59. Savoca, V. et al. TERRA stability is regulated by RALY and polyadenylation in a telomere-specific manner. Cell Rep. 42, 112406 (2023).

60. Gilot, D. et al. A non-coding function of TYRP1 mRNA promotes melanoma growth. Nat. Cell Biol. 19, 1348–1357 (2017).

61. Migault, M., Donnou-Fournet, E., Galibert, M.-D. & Gilot, D. Definition and identification of small RNA sponges: Focus on miRNA sequestration. Methods 117, 35–47 (2017).

62. Hagedorn, P. H. et al. Locked nucleic acid: modality, diversity, and drug discovery. Drug Discov. Today 23, 101–114 (2018).

63. Renneberg, D., Bouliong, E., Reber, U., Schümperli, D. & Leumann, C. J. Antisense properties of tricyclo-DNA. Nucleic Acids Res. 30, 2751–2757 (2002).

64. Goyenvalle, A. et al. Functional correction in mouse models of muscular dystrophy using exon-skipping tricyclo-DNA oligomers. Nat. Med. 21, 270–275 (2015).

65. Mitic, N. et al. Dissecting the spatiotemporal diversity of adult neural stem cells. Mol. Syst. Biol. 20, 321–337 (2024).

66. Liberzon, A. et al. Molecular signatures database (MSigDB) 3.0. Bioinformatics 27, 1739–1740 (2011).

67. Zhang Hao. zhanghao-njmu/SCP. (2026).

