## Supplementary Figures for "TERRA engages MDA5 to shape the immune landscape in ALT brain tumors"

SUPPLEMENTARY FIGURE 1 See comments for fig. 1A

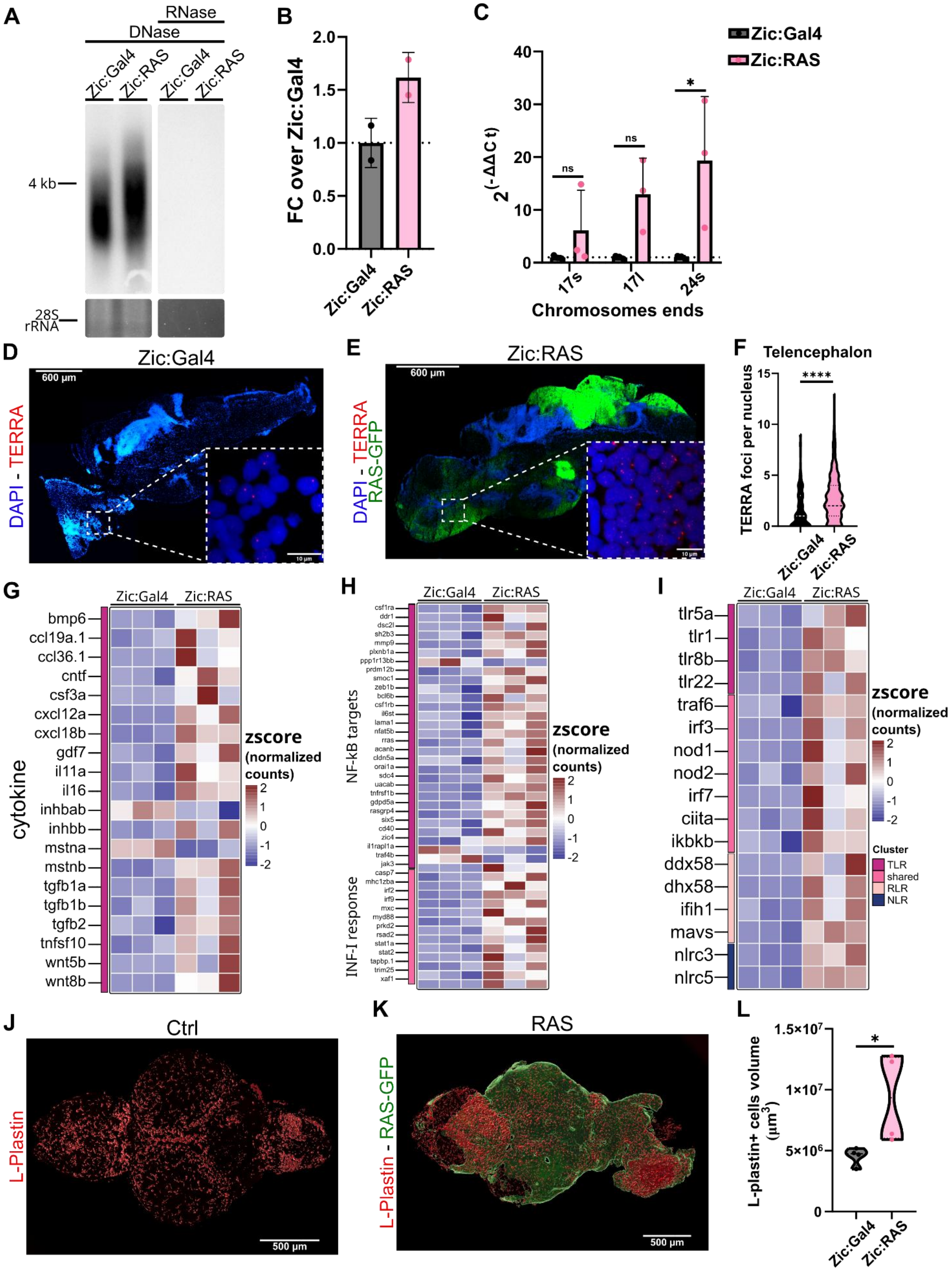

**Supplementary Figure 1: Increase of TERRA expression in ALT+ brain tumors correlates with increased inflammatory responses**

(A) Representative Northern blot showing TERRA levels in Zic:Gal4 (control) versus Zic:RAS adult brain tissue (loading control shown below).

(B) Quantification of relative signal intensity normalized over control.

(C) qPCR analysis of TERRA expressed from telomeres 17s, 17l and 24s in adult brain tissue. Quantity is expressed as fold change of Zic:RAS over Zic:Gal4, normalized over HK gene. (Statistics: 2way Anova with Sidak's multiple comparisons test , n=3, *p<=0.05, $^{ns}$p>0.05).

(D, E) RNA fish performed on Zic:Gal4 (D) or Zic:RAS (E) adult brains, highlighting cell nuclei (blue), hRAS-GFP (green) expression and TERRA foci (red).

(F) Quantification of TERRA foci per nucleus in the cerebellum of Zic:Gal4 and Zic:RAS adult fish (Statistics: Mann-Whitney test, Zic:Gal4 n=184, Zic:RAS n=247, *p<=0.05).

(G, H, I) Heatmap showing the z-scores of the expression of cytokines (G), NF-kB targets, INF-I response (H) and RNA-sensing genes (I) in three brains of Zic:Gal4 or Zic:RAS adult fish.

(J, K) Representative Fiji-processed max-projection WM-IF images of adult brains of Zic:Gal4 (J) and Zic:RAS (K), showing L-plastin staining (red) and hRAS-GFP expression (green). Scale bar: 200 µm.

(L) Quantification of the total volume µm$^3$ occupied by L-plastin+ per brain. (Statistics: unpaired T-test, n=4, *p<=0.05).

### SUPPLEMENTARY FIGURE 2

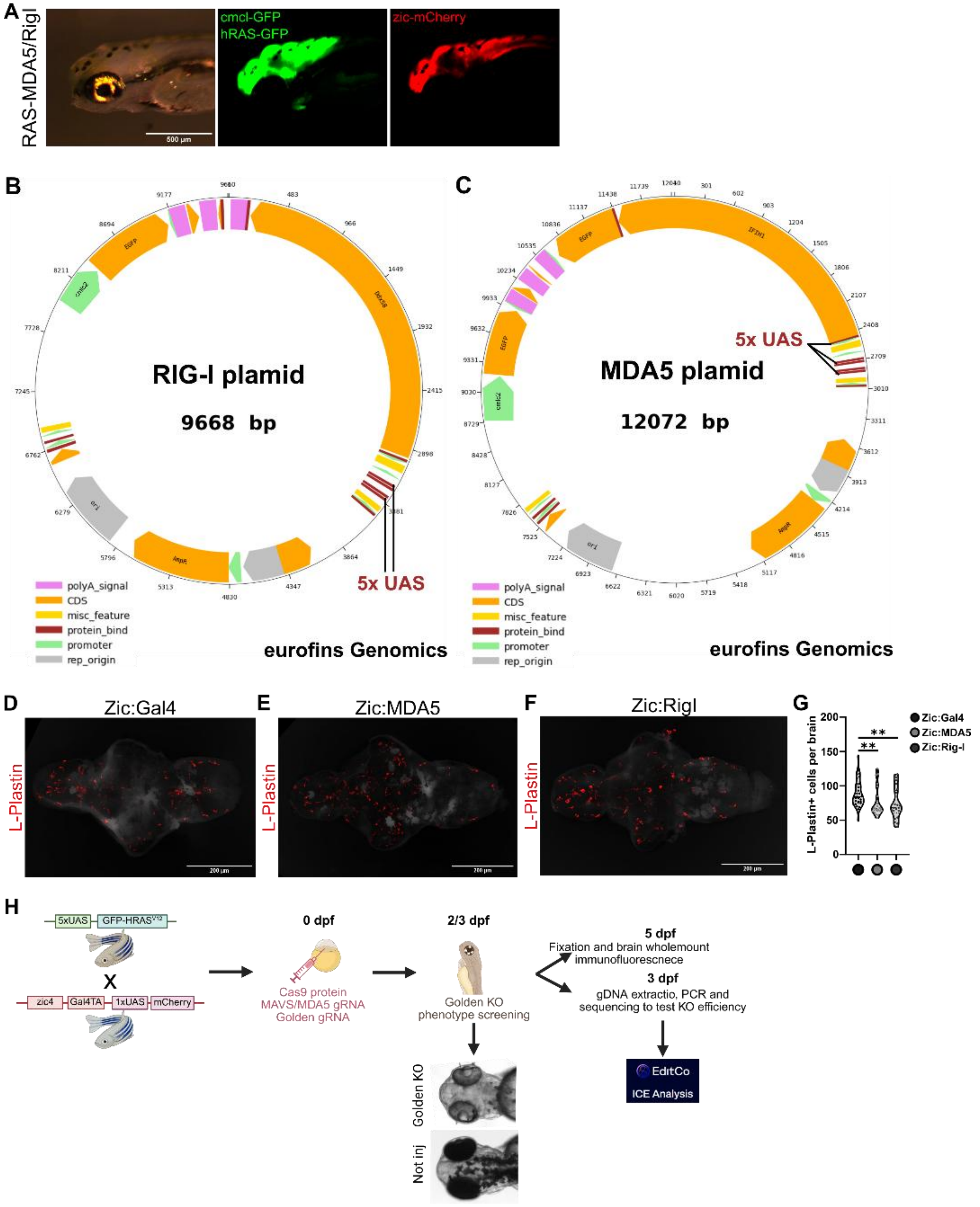

**Supplementary figure 2: Genetic manipulation of the RLR pathway alters immune infiltration in Zic:RAS ALT+ tumors.**

(A) Images of a double-transgenic zebrafish embryo (RAS-MDA5/Rig-I). Panels show GFP fluorescence (indicating cmlc-GFP / hRAS-GFP expression) and mCherry fluorescence. Scale bar: 500μm.

(B, C) Plasmid maps for the expression vectors encoding (B) mouse Rig-I (9668 bp) and (C) human MDA5 (12072 bp), showing the 5x UAS driver elements, the cmcl-GFP reporter and Ddx58 (Rig-I) or IFIH1 (MDA5) coding sequence respectively. Sequence and images obtained through Eurofins Genomics.

(D-F) Representative images of WM-IF on the brain of 5 dpf larvae for L-plastin performed on Zic:Gal4, Zic:MDA5 and Zic:RIG-I animals. L-plastin cells are in red. Images are processed with Fiji. Scale bar: 200μm.

(G) Quantification of the number of L-plastin cells (statistics: Dunn's multiple comparisons test, Zic:Gal4 n=34, Zic:MDA5 n=25, Zic:RIG-I n=26; **p<0.01).

(H) Schematic representation of the protocol used to obtain and analyze crispant larvae (created with Biorender.com).

**SUPPLEMENTARY FIGURE 3**

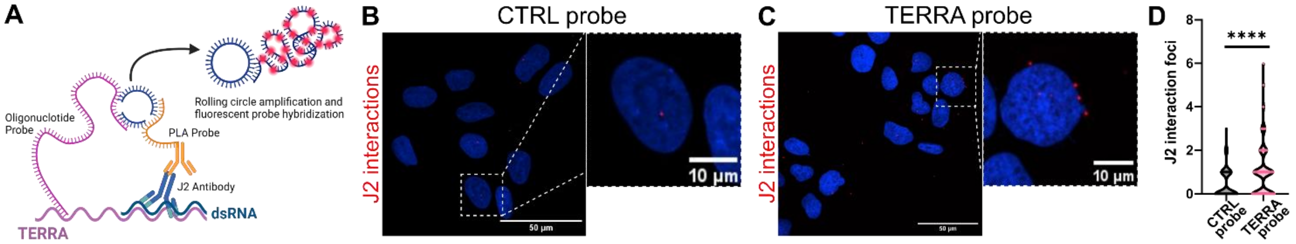


**Supplementary figure 3: Increased TERRA engages the RLR pathway to induce inflammation**

(A) Schematic representation of PLA technology used to detect the ds-TERRA fraction using the J2 antibody, specific for dsRNA (Created with Biorender.com)

(B-C) Representative images of PLA performed in U2OS cells with J2 antibody and CTRL (B) or TERRA (C) probe. TERRA interaction foci are in red, DAPI is in blue. Images are processed with Fiji. Scale bar: 50 µm and 10 µm.

(D) Relative quantification of the J2 interaction foci in U2OS (Statistics: Mann Whitney test; CTRL probe n=99, TERRA probe n=167; ****p<0.0001).

**SUPPLEMENTARY FIGURE 4**

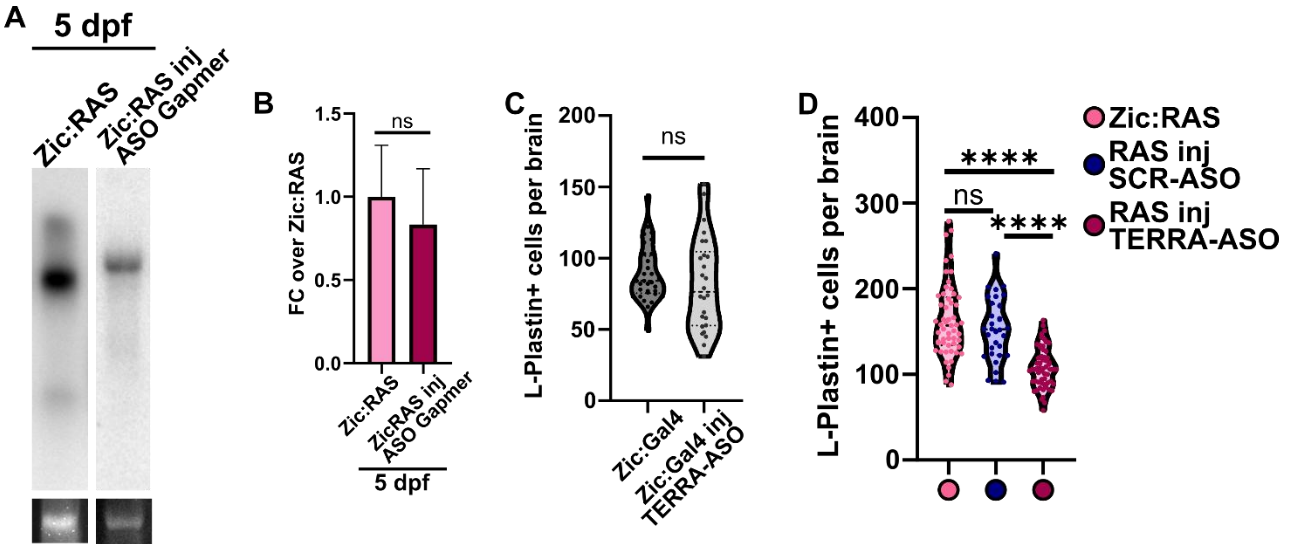


**Supplementary figure 4: Antisense TERRA reduces inflammation, tumor cell proliferation and prolong survival of tumor bearing larvae**

(A) Representative Northern blot (or RNA gel) analysis showing TERRA levels in Zic:Gal4 (control) versus Zic:RAS 5 dpf larvae (loading control shown below).

(B) Quantification of relative signal intensity normalized over control. (Statistics: unpaired T-test, Zic:RAS n=5; ZicRAS inj ASO Gapmer n=3; nsp>0.05).

(C) Quantification of the number of L-plastin cells per brain (Statistics: Mann Whitney test, Zic:Gal4 n=33; Zic:Gal4 inj TERRA-ASO n=26; nsp>0.05).

(D) Quantification of the number of L-plastin cells per brain (Statistics: Kruskal-Wallis test with Dunn's multiple comparisons test; Zic:RAS n=55; RAS inj TERRA-ASO n=59; RAS inj SCR-ASO n=32; nsp>0.05).

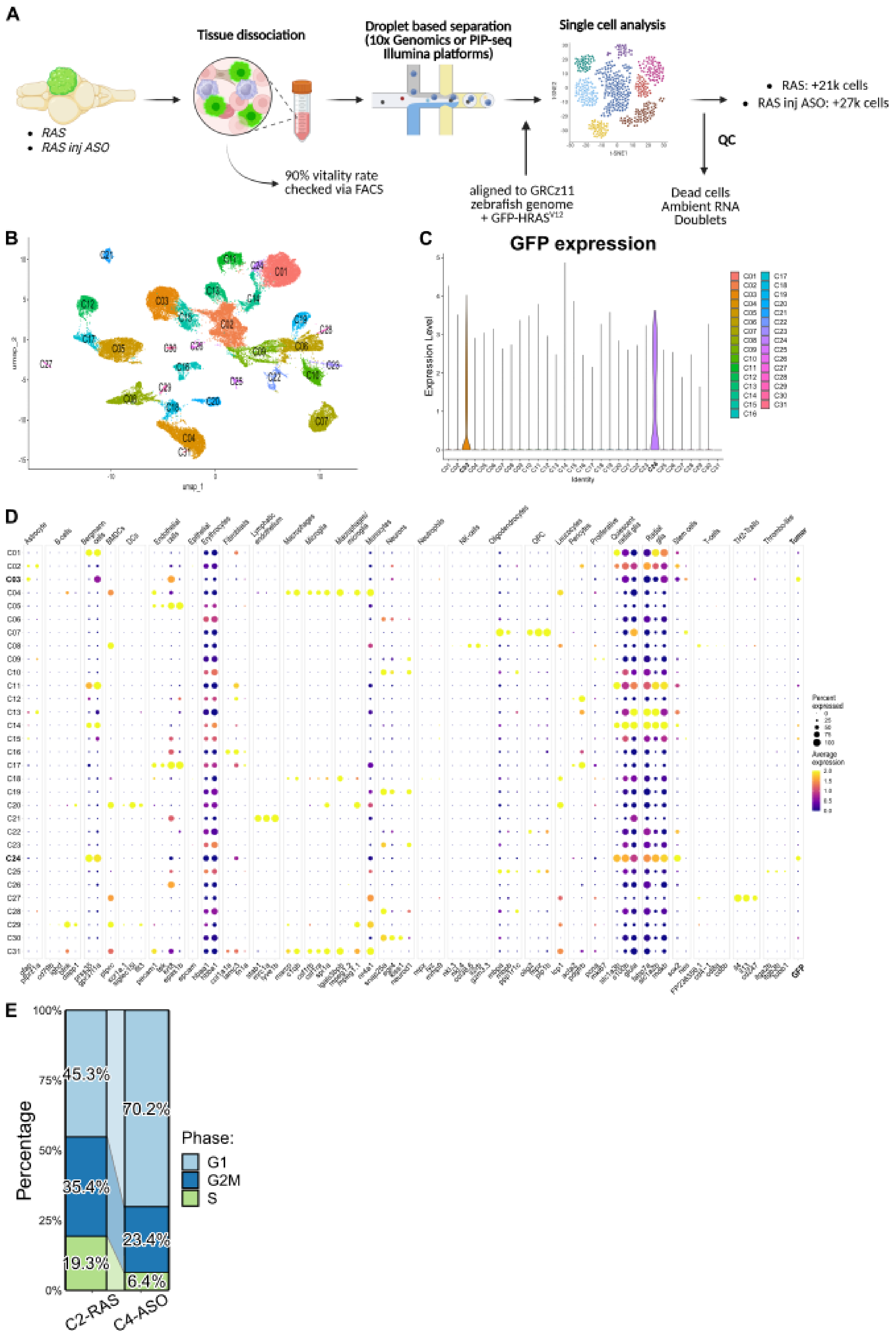


SUPPLEMENTARY FIGURE 5

**Supplementary figure 5: TERRA ASO treatment affect ALT+ cancer cells in a long-lasting way**

(A) scRNA-seq experimental workflow: Brain tissues from Zic:RAS and Zic:RAS-ASO were dissociated (>90% viability) and processed via droplet-based scRNA-seq technologies (10x Genomics/PIP-seq). Reads were aligned to the GRCz11 genome supplemented with transgene GFP-hRASV12. After filtering we obtained more than 48k cells.

(B) UMAP of 49279 aggregated single cells coming from the two models showing the different clusters obtained with unsupervised clustering.

(C) Violin plot showing the expression of the transgene GFP-hRASV12 in the 31 clusters found in the dataset. Colors indicating the different clusters.

(D) Dotplot showing the expression of the genes used for the manual annotation of the 31 clusters found in the dataset via unsupervised clustering. Dot size indicates the percentage of cells in the cluster expressing the gene and the color the average expression (high expression in yellow and low expression in blue)

(E) Stacked barplot showing the percentage of cells in the G1,G2, or S cell cycle phase in the C2-RAS and C4-RAS clusters
